# Generalized Leaky Integrate-And-Fire Models Classify Multiple Neuron Types

**DOI:** 10.1101/104703

**Authors:** Corinne Teeter, Ramakrishnan Iyer, Vilas Menon, Nathan Gouwens, David Feng, Jim Berg, Nicholas Cain, Christof Koch, Stefan Mihalas

## Abstract

In the mammalian neocortex, there is a high diversity of neuronal types. To facilitate construction of system models with multiple cell types, we generate a database of point models associated with the Allen Cell Types Database. We construct a series of generalized integrate-and-fire (GLIF) models of increasing complexity aiming to reproduce the spiking behaviors of the recorded neurons. We test the performance of these GLIF models on data from 771 neurons from 14 transgenic lines, with increasing model performance for more complex models. To answer how complex a model needs to be to reproduce the number of electrophysiological cell types, we perform unsupervised clustering on the parameters extracted from these models. The number of clusters is smaller for individual model types, but when combining all GLIF parameters 18 clusters are obtained, while 11 clusters are obtained using 16 electrophysiological features. Therefore, these low dimensional models have the capacity to characterize the diversity of cell types without the need for a priori defined features.

## 1 Introduction

The problem of understanding the complexity of the brain has been central to neuroscience. Cell types are a conceptual simplification intended to reduce this complexity. A large scale effort at the Allen Institute for Brain Science has focused on characterizing the diversity of cell types in primary visual cortex of the adult mouse using electrophysiology, morphological reconstructions, connectivity, and modeling in one standardized effort. This has resulted in the Allen Cell Type Database that is publicly and freely available at http://celltypes.brain-map.org. It includes both morphological and electrophysiological data of genetically identi ed neurons mapped to the common coordinate framework. Morphologically and biophysically detailed as well a simple generalized linear integrate-and-fire point neuron models have been generated (Allen Institute for Brain Science, 2016a) to reproduce the highly standardized data.

In this study, we address the problem of cell types using point models to provide a dimensionality reduction to the most relevant aspects of neuronal input/output transformations. We show the ability of simple generalized leaky integrate and fire point models to both reproduce the spike times of biological data and effectively reduce the biological space to a set of useful parameters. Beyond the benefits of clarifying mechanisms for single neuron behavior, single neuron models are used in larger network models that attempt to explain network computation. Associated with this study we provide a large scale database of point neuron models.

Many neuron models have been developed to describe and recreate various aspects of neuronal behavior (Koch, 2004). At one end of the spectrum are the morphologically and biophysically realistic Hodgkin-Huxley like models (Hodgkin and Huxley, 1952; Hay et al., 2011). Their strength relies in their capacity to map between multiple observables: morphology, electrophysiology, intra-cellular calcium concentration and levels of expression and patterns of distributions of ionic currents. At the other end are highly simplified models, in particular the integrate-and-fire (IF) family of models (Lapicque, 1907). Although adding complexity to a model may increase the ability of that model to recreate certain behavior,finding the right parameters for complex models becomes a challenge (Prinz et al., 2003), and the computational power needed to simulate sophisticated neural models can be quite large (Markram et al., 2015). Therefore, ideally one would use a computationally minimal model adequate to recreate and to understand the desired behavior (Gollisch and Meister, 2010). In order to both facilitate network modeling efforts and understand the complexity of cell types, models many levels of sophistication have been generated (Allen Institute for Brain Science, 2016a) to recreate biological data. However, the level of complexity needed to meaningfully describe biological data is not understood.

One simplification which significantly reduces model complexity is to represent the entire dendritic tree, soma and axonal hillock by a single compartment, while maintaining the dynamics of the individual conductances (Koch, 2004). This approximation is especially warranted when characterizing neurons via somatic current injection and voltage recording.

Several models can reproduce a wide variety of spiking behavior, while reducing the number of dimensions to two (FitzHugh, 1961; Morris and Lecar, 1981). In 2003, Izhikevich also created a simple model that recreated several stereotypical realistic firing patterns by reducing the dimensionality of the Hodgkin-Huxley model (Izhikevich et al., 2003; Izhikevich, 2007). Although low-dimensional (Izhikevich, 2004), this model is non-linear which creates difficulties optimizing the model parameters to achieve biological spiking behavior. Similar firing patterns can be obtained using the adaptive exponential (AdEx) neuronal model (Brette and Gerstner, 2005).

Threshold-based models, which represent the spike initiation as a threshold rather than a dynamical system, are based of the simple leaky integrate and fire (LIF) (Lapicque, 1907) model. Their parameters can more easily be fit to biological data (Paninski et al., 2004; Rossant et al., 2010; Mihalas and Niebur, 2009; Dong et al., 2011; Mensi et al., 2012; Pozzorini et al., 2015). For a more in-depth characterization of the diversity of neuron models, see the review (Herz et al., 2006), and for their capacity to reproduce spike times see (Gerstner and Naud, 2009).

Here, we use the leaky integrate-and-fire (LIF) framework, due to the versatility of adding multiple mechanisms and the capacity to rapidly find unique parameters for them. A standard LIF model was the starting point, progressing to more generalized leaky integrate-and-fire (GLIF) models schematized in Figure 2a. In these models, the mechanisms are separated by time scale: none of the fast processes associated with the action potential itself are included directly in the dynamics. Some of these are, however, accounted for in the reset rules which map the state before the spike to the state after (i.e. voltage and threshold). Unlike the simple LIF which assumes all spikes are stereotyped and reset to the same state, the generalized reset rule (R) assumes that the spikes are sufficiently similar such that a mapping between the state before and after the spike can be found. In this study, we search only for linear mappings. The rapid changes of the action potentials are followed by slower dynamics which affect a neuron’s state: e.g. slow inactivation of voltage-dependent currents which activate during the spike, or *Ca*^2+^-dependent potassium currents. We assume that they have a stereotyped activation following a spike, and we bundle all these effects into a set of afterspike currents at different time scales (ASC), and into a change of the instantaneous threshold. Finally, slow depolarization can lead to partial inactivation of the voltage-dependent sodium current which generates a spike. We incorporate this mechanism into a change of the threshold which is dependent on the membrane potential (AT). The GLIF framework (Mihalas and Niebur, 2009) is sufficiently general such that multiple additional mechanisms can be added. How such a framework can be used to rapidly fit neuronal models is known (Pozzorini et al., 2015). The current work uses similar methods to (Pozzorini et al., 2015), with a set of differences: 1. More complex reset rules are used which map the state of the neuron before the spike to that after. These are essential for differentiating between neuron types. 2. We include a membrane potential dependent adaptation of threshold. 3. Exponential bases functions are used for the time dependencies. These allow a much sparser representation which is essential for clustering. 4. We use a direct measurement of noise in the membrane potential of a neuron rather than a probabilistic threshold. 5. The variance in the input current is smaller, closer to that observed in vivo.

When classifying neuron types based on a selection of features, one question which is difficult to address is the importance of these features to the function of the circuit. By doing the clustering on parameters which link somatic input of currents to spike train, we construct classes which are relevant for simulations where spike times are a desired observable. One of the advantages of GLIF neurons is their capacity to obtain unique parameters for diverse mechanisms (Mensi et al., 2012; Dong et al., 2013), which we show here, allow for classiffications on model parameters.

Here we demonstrate the great power of computationally simple GLIF models to both reproduce the spike times of biological data and reduce the vast biological parameter space to a low dimensional space which can classify neurons with a specificity equal to that of common eletrophysiological features. We characterize the parameters of the GLIF models from the different transgenic lines and illustrate the neuron that best recreates the biological spike train for each of the transgenic lines. In addition, we characterize common electrophysiological features associated with the different GLIF clusters.

## Results

### 2.1 Model Definitions

We developed five different GLIF models, each of which successively incorporates the time evolution of one to five state variables. The set of five state variables X = {*V*(t), Θ_s_(t),*I*_(j=1,2)_(t), Θ_V_(t)} represent the neuronal membrane potential, the spike-dependent component of the threshold, two after-spike currents and the membrane potential-dependent component of the threshold respectively. Figure 1 illustrates the iteration between the state variables in the model equations. It is assumed that these state variables evolve in a linear manner between spikes:

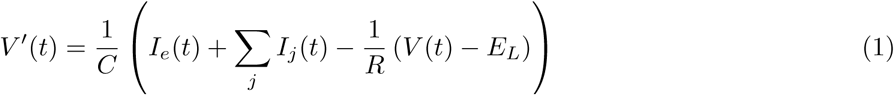

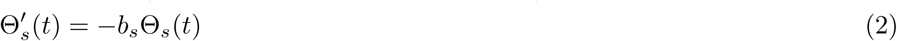

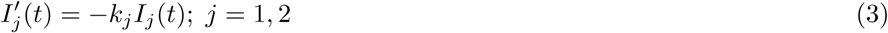

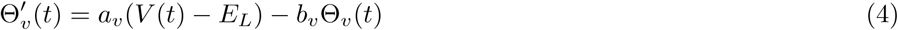

where *C* represents the neurons’s capacitance, *R*, the membrane resistance, *E*_*L*_, the resting membrane potential, *I*_e_, the external current, 1/*k*_*j*_, 1/*b*_*s*_, 1/*b*_*v*_ are the time constant of the afterspike currents, spike and voltage dependence of the threshold and *a*_*v*_ couples the membrane potential to the threshold.

**Figure 1:**
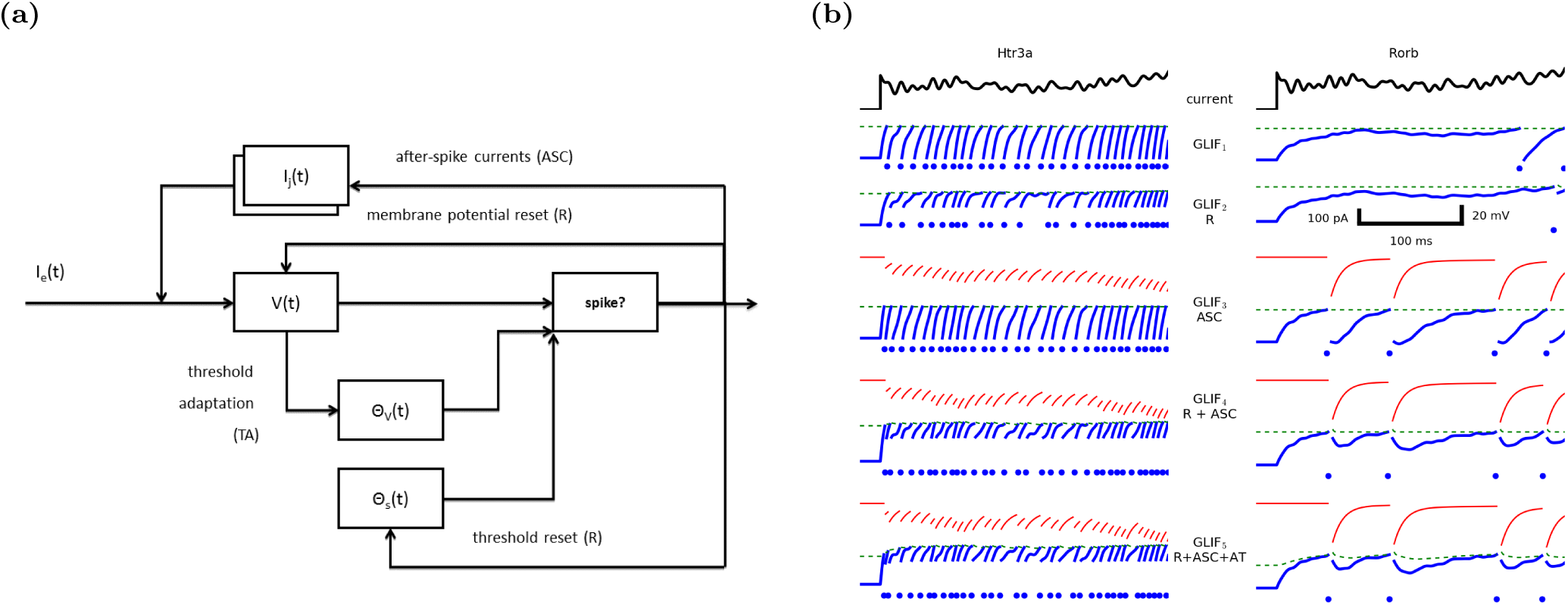
(a) Schematic describing the mechanisms included in our family of generalized leaky integrate-and-fire (GLIF) models used to describe the evolution of the trans-membrane potential and spiking of close to a thousand neurons recorded under highly standardized conditions. The input is a current, *I_e_*(*t*), injected via the patch electrode. The output is the membrane potential, *V*(*t*). 1D model: *GLIF*_1_ model contains one variable, *V*(*t*), which is reset when it reaches a fixed threshold. 2D model: *GLIF*_2_ models include a second variable, spike induced threshold Θ_*S*_(*t*), which dynamically defines the value of the threshold. Both *V*(*t*) and Θ_S_(*t*) after the spike linearly depend on their values before the spike; between spikes their evolution equations are independent. 3D model: *GLIF*_3_ include *V*(*t*) and two variables corresponding to two after spike currents with different time constants. 4D model: *GLIF*_4_ combines the spike induced threshold changes and after spike currents. 5D model: *GLIF*_5_ adds Θ_*V*_(*t*) as a membrane potential dependent component of the threshold. (b) Illustration of the five distinct GLIF models using exemplary recorded neurons from two different transgenic lines (left and right column). The injected current, *I*_e_(*t*), is plotted at the top in black. In all panels, the model trans-membrane potential, *V*(*t*), is pictured in blue (spike rasters are marked below), and the threshold, in dashed green. For models that contain afterspike currents, the total current is in red. The scale bar applies to all panels except to the injected current. *GLIF*_1_: Equivalent to the standard LIF model, when *V*(*t*) reaches the constant threshold, a spike is produced. After a short refractory period, *V* is reset to a constant value. *GLIF*_2_: The threshold Θ_*s*_ modestly increase with each spike, and decays to a constant. When *V*(*t*) reaches Θ_*s*_, the neuron spikes. After the refractory period, both V and are reset to a value which is dependent on the state of the neuron before the spike. *GLIF*_3_: Every action potential induces afterspike currents which decay back to zero. The level of induction depends on the state of the neuron before the spike. *GLIF*_4_: Here, spike-induced changes in threshold and afterspike currents are combined and reset for all variables which are dependent on the state of the neuron before the spike. *GLIF*_5_: The addition of a voltage-dependence to the threshold Θ.

If *V*(*t*) > Θ_*v*_(*t*) + Θ_*s*_(*t*) + Θ_∞_, a spike is generated and the state variables are updated with a linear dependence on the state before the spike:

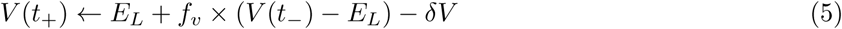

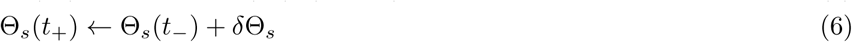

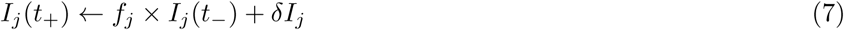

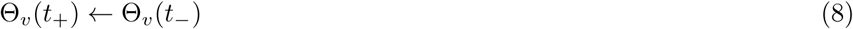

The 1D model, *GLIF*_1_ is represented by equation 1, with a simple reset. *GLIF*_2_ adds Θ_*S*_ and the more complex reset rules (equations 5 and 6). *GLIF*_3_ uses membrane potential and two spike induced currents, *GLIF*_4_ uses membrane potential, spike induced threshold dependence and afterspike currents, while GLIF_5_ uses all the variables described. A detailed description of the models along with their mathematical equations are available in the “4.1 Model Deffinitions” section of the Supplementary Material.

### 2.2 Data

Intracellular electrophysiological recordings were carried out via a highly standardized process (Allen Institute for Brain Science, 2016b). The majority of the data (approx. 600 cells) can be accessed on the Allen Cell Types Database. Data that pass standardized electrophysiology quality control criteria but fail downstream quality control criteria such as imaging or reconstruction are included in this manuscript but are not released in the Allen Cell Types Database. The available data consists of *in vitro* electrophysiology data collected from 19 different transgenic lines, recording from both labeled (Cre positive) and non labeled (Cre negative) neurons (Figure 2).

**Figure 2:**
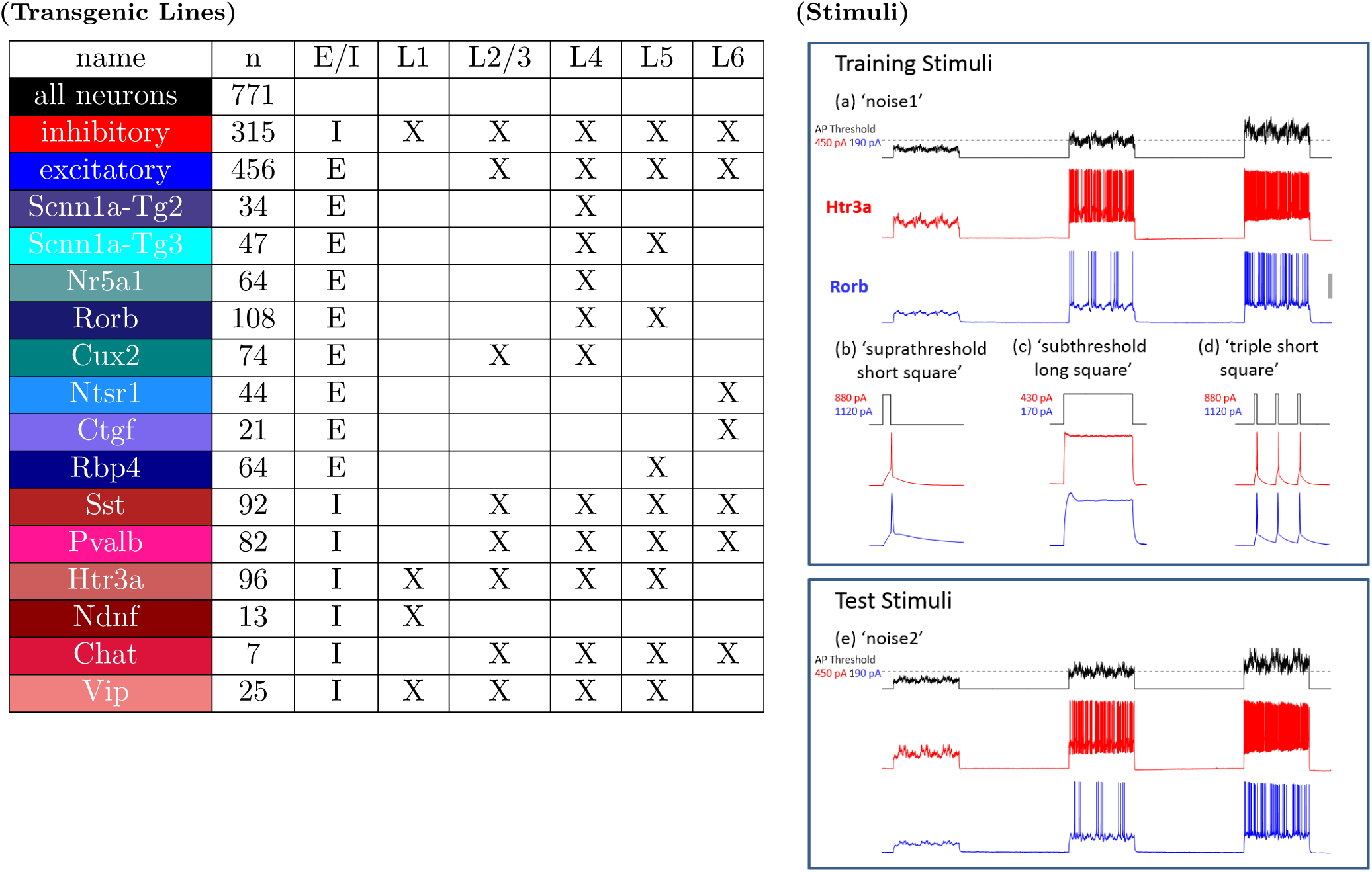
771 different neurons from 14 transgenic lines containing all the required stimuli on the Allen Cell Types Database are considered in this study. (left) Illustrated colors correspond to the different transgenic lines in all figures.’n’ describes the number of neurons for which the lowest level model (*GLIF*_1_) could be generated. Transgenic lines can identify either inhibitory (I) or excitatory (E) cells which reside in layer 1 (L1) through layer 6 (L6). (right) A minimal set of stimuli were required for training and testing di erent GLIF models. GLIF models were trained using (a) at least two repeats of pink noise stimuli (3 s each, 1/f distribution of power, 1 - 100 Hz) with amplitudes centered at 75, 100, and 125 percent of action potential threshold, (b) a short (3 ms) just suprathreshold pulse in order to fit the instantaneous threshold, Θ_∞_, (c) a long square (1 s) pulse just below threshold to estimate the intrinsic noise present in the voltage traces (used in the post hoc optimization step), and (d) a series of three peri-threshold short pulse sets for any model with reset rules (*GLIF*_2_, *GLIF*_4_, and *GLIF*_5_). GLIF models were then tested using a holdout stimulus set (b) of at least two sweeps of a second pink noise stimuli generated in an identical manner to the training but initialized with a different random seed. Representative data shown from an Htr3a-Cre positive neuron (resting membrane potential (RMP) −74 mV) and a Rorb-Cre positive neuron (RMP = −77 mV). Scale bar in (a) corresponds to 40 mV (panels a, b, d, e) or 15 mV (panel c).

Transgenic lines selective for different types of neurons were used for targeting. Excitatory neurons from each layer are identified using layer-selective lines (Figure 2). For inhibitory neurons, cells are targeted across lamina using lines selective for known neuronal subtypes Tasic et al. (2016). Cells from each of the three major inhibitory subtypes (Pvalb (primarily basket cells), Sst (primarily Martinotti cells), and Htr3a (diverse morphologies)) are represented. The Vip, Ndnf, and Chat Cre lines label a subtype of Htr3a-positive interneurons.

Only transgenic lines which contained more than 7 neurons that had the necessary data to create a basic *GLIF*_1_ model described in Figure 2 were included in this study. In addition, models which had biologically unrealistic parameter values are eliminated from the data set (Please see section “4.7 Exclusion Criteria” in the Supplementary Material for a detailed explanation of excluded data). Figure 2 summarizes the properties of the different transgenic lines considered here. Colors and short hand names represent all transgenic line data throughout this manuscript. In general, shades of red represent inhibitory transgenic lines and shades of blue represent excitatory lines. Overall a total of 771 neurons from 14 transgenic lines meet the criteria for fitting the *GLIF*_1_ and *GLIF*_3_ models. Of these, 289 neurons meet the additional requirements for models with reset rules (*GLIF*_2_ and *GLIF*_4_) and 288 neurons meet all the criteria for a *GIF*_5_ model.

Throughout the manuscript data and models from two exemplary canonical neurons are consistently shown as examples: an Htr3 inhibitory neuron (specimen ID 474637203) and a Rorb pyramidal neuron (specimen ID 479766169).

We designed a set of stimuli optimized for testing parameters of our GLIF models (Figure 2). The noise stimuli were picked to have similar statistics to *in vivo* patch clamp recordings. Using stimuli with highly varied structure is important, as it allows the model to explore the parameter space. Using stereotyped stimuli, e.g. square pulses, would present only a small subset of possible histories (both spiking and sub-threshold), and bias the dataset to a non-physiological regime. A short (1s) stimulus chunk is repeated three times to allow the analysis of short term adaptation. These three second bouts are repeated at different amplitudes, one sub-threshold, one peri-threshold and one supra-threshold to explore di erent parameter regimes. This stimulus sequence is repeated at least twice to check for reliability of the neurons under the same stimulus history. A second instantiation of a noise with the same characteristics is used to check the performance of the model under naturalistic stimulus conditions. A very short (3 ms) square stimulus of multiple amplitudes is used to compute the neuron’s instantaneous threshold. A series of such short squares at different frequencies are used to characterize the spike-induced changes in the absence of additional membrane potential-induced changes. These are part of the electrophysiological stimulation protocol which is the subject of a different study.

### 2.3 Post-hoc optimization of instantaneous threshold, Θ_∞_

GLIF model fitting is achieved in two steps. In the first step, parameters are fit directly from the electrophysiological data using linear methods described in detail in section “4.2 Parameter fitting and distributions” of the Supplementary Material: *E_L_* from the mean membrane potential at rest, *R*, and *C* from fitting the membrane potential during a subthreshold noise stimulus, *δI_j_*; *k_j_* from fitting the membrane potential between spikes during a suprathreshold stimulus, *f_v_*; *δt*, *δV* relating the potential before and after a spike during during the noise stimulus, *b_s_*; *δ*Θ_*S*_ from the potential of subsequent spikes during a series of brief square pulses, *a_v_* and *b_v_* from the potential of subsequent spikes during the suprathreshold noise stimulus. Slices from this parameter space are shown in Figure 3.

**Figure 3:**
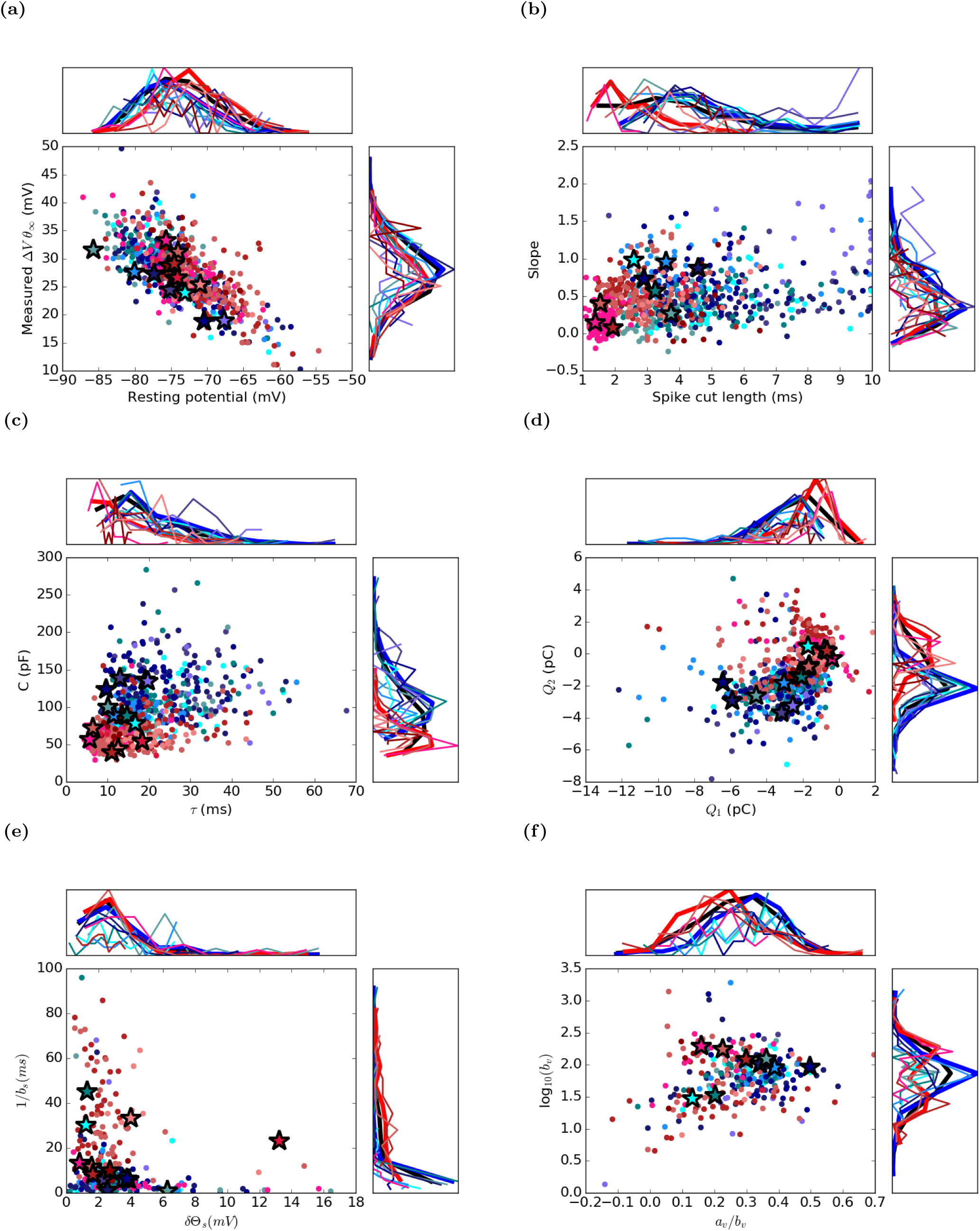
Slices from the complex parameter space fit from electrophysiological data. Five parameters are fit to the basic *GLIF*_1_ (or LIF) model. Four additional parameters are fit to models with after-spike currents (ASC) (*GLIF*_3_, *GLIF*_4_, *GLIF*_5_) and another two parameters are fit to models with reset rules (R) (*GLIF*_2_, *GLIF*_4_, *GLIF*_5_). Finally two additional parameters are fit for the *GLIF*_5_ model containing adapting threshold (AT), resulting in a total of 15 parameters fit for the most sophisticated model considered here (Table 1). Detailed parameter fitting methodology (Figure 2) is available in the Supplementary Material. (a) Resting potential is measured as the average voltage during rest before training noise (noise1) current is injected. The threshold relative to rest, Δ*V*, is measured by subtracting resting voltage from the threshold obtained from the suprathreshold short square pulse. (b) The spike waveform is removed from the voltage trace by aligning all spikes and fitting a line to the voltage before and after a spike. The best fit line within a window of 10 ms after spike initiation was chosen (Figure 8). This spike width is used in all models, and voltage measurements before and after a spike are used to reset voltage in (*GLIF*_2_, *GLIF*_4_, and *GLIF*_5_). (c) Capacitance and resistance are fit via linear regression to subthreshold voltage (Figure 9). The membrane time constant, *τ* = RC is plotted. (d) Total charges of the fast Q_1_ and slow Q_2_ after-spike currents deposited each time there is a spike. (e) The amplitude, *a_s_*, and decay *b_s_* of the spiking component of the threshold, Δ_s_, (used in *GLIF*_2_, *GLIF*_4_, and *GLIF*_5_) is t to the triple short square data set (Figure 10). (f) In the *GLIF*_5_, the threshold is influenced by the voltage of the neuron according to (Equation 4). The two parameters of this equation are plotted here.

In the second step, the instantaneous threshold, Θ_∞_, was optimized to fit the probability of a neuron model reproducing observed spike times. This is realized using a non-linear Nelder Mead optimization strategy as described in detail in section “4.4 Post-Hoc Optimization” of the Supplementary Material. Θ_∞_ was chosen for further optimization because it is an essential parameter in the overall excitability of neurons and is likely the best parameter to counteract any error in the fitting or the inherent error introduced by simplifying any complex system to a simple model. Although Θ_∞_ did change between the fitting and post-hoc optimization step, on average the value remains consistent (See Figure 7 c and d in the Supplementary Material).

### 2.4 Model performance

After optimization, we quantified the model performance by how much of the temporal variance in the spike times of the data in response to a *in vivo* like stimulus can be explained by the model at a 10 ms temporal scale. Please see section 4.5 in the Supplementary Material. *GLIF*_1_ (equivalent to a leaky integrate and fire model) has a surprisingly high explained variance of 70%. It is better for inhibitory than for excitatory neurons: 76% versus 67% (Figure 5, Table 4). The addition of reset rules and after spike currents independently do not significantly improve the explained variance. However, when both are jointly added, they improve the explained variance from 70% to 74% (86% for inhibitory and 71% for excitatory neurons). On top of that, the addition of the adaptive threshold provides a significant but smaller improvement from 74% to 76% (87% for inhibitory and 73% for excitatory neurons). Performance of individual transgenic lines can be viewed in the Supplementary Material (Figures 13 and 14)

**Figure 4:**
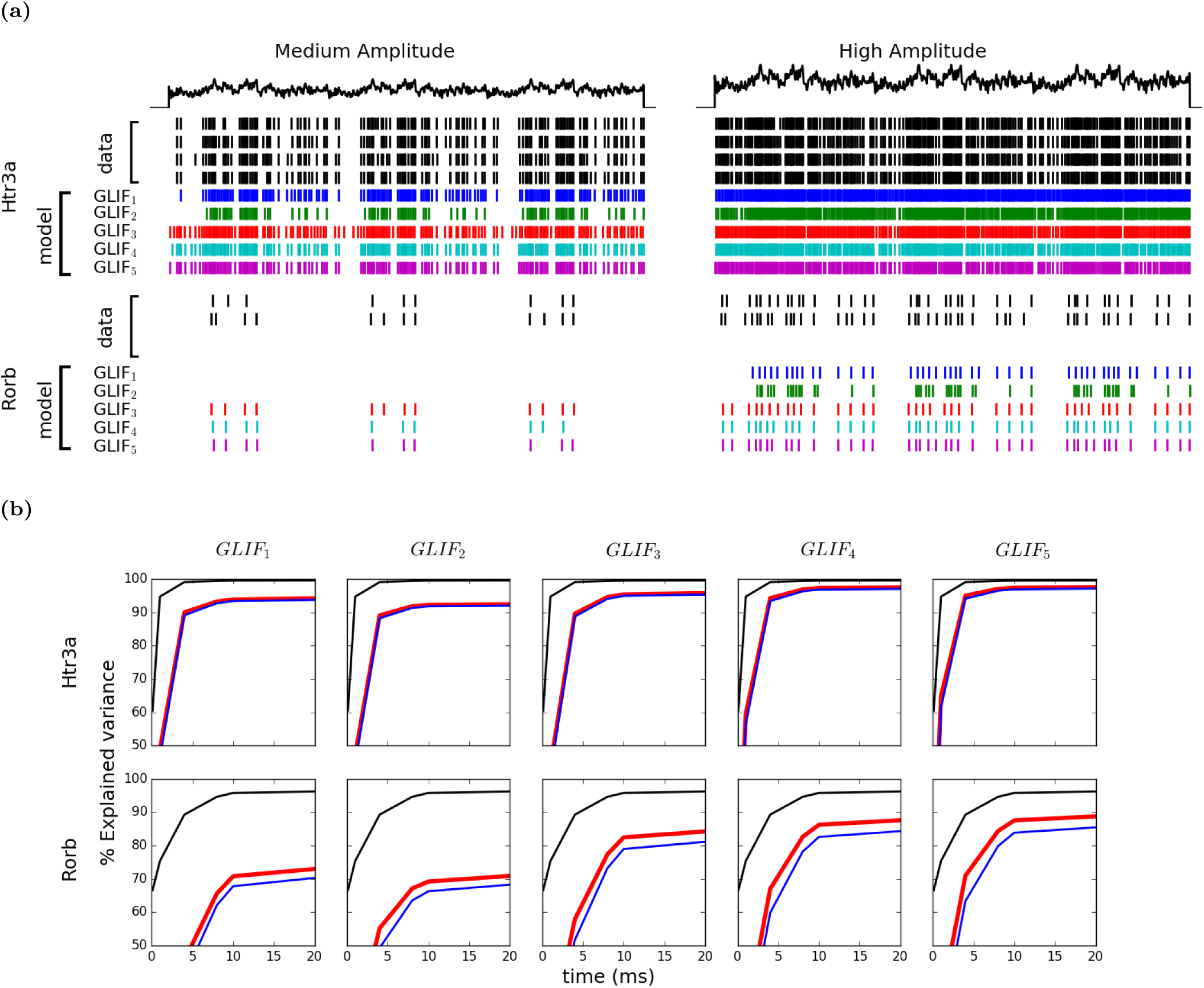
Rastergrams and explained variance of biological data and all optimized model levels for ‘hold out’ test data. (a) Data for the two example cells: Htr3a 477975366 and Rorb 314822529. Injected current shown in black. Black rasters are spikes from recorded neurons to repeated current injections. Colored rasters correspond to the five different, deterministic models. The current injection is 3 s long. It is observed that as the *GLIF*_1_ and *GLIF*_2_ do not have a spike frequency adaptation mechanism, they have trouble reproducing simultaneously the ring patterns at multiple input amplitudes. (b) Explained variance for the different model levels at different levels of time window resolution *Δt*. The black lines represent the explained variance of the data (how well the neuron repeats its own spiking behavior). This trace would reach 100% if the spike times of the data were all exactly at the same time in each repeated stimulus: the fact that they are not 100% reects the intrinsic variation in the spike times within the experimental data. The blue line illustrates the pairwise explained variance of the model with the data. Because the model can not be expected to explain the data better than the data can explain itself, the red line is the ratio of the pairwise explained variance of the model (blue) and the data divided by the explained variance of the data (black). This ratio value at a Δ t=10 ms time bin is used for the explained variance performance metric in the main text.

**Figure 5:**
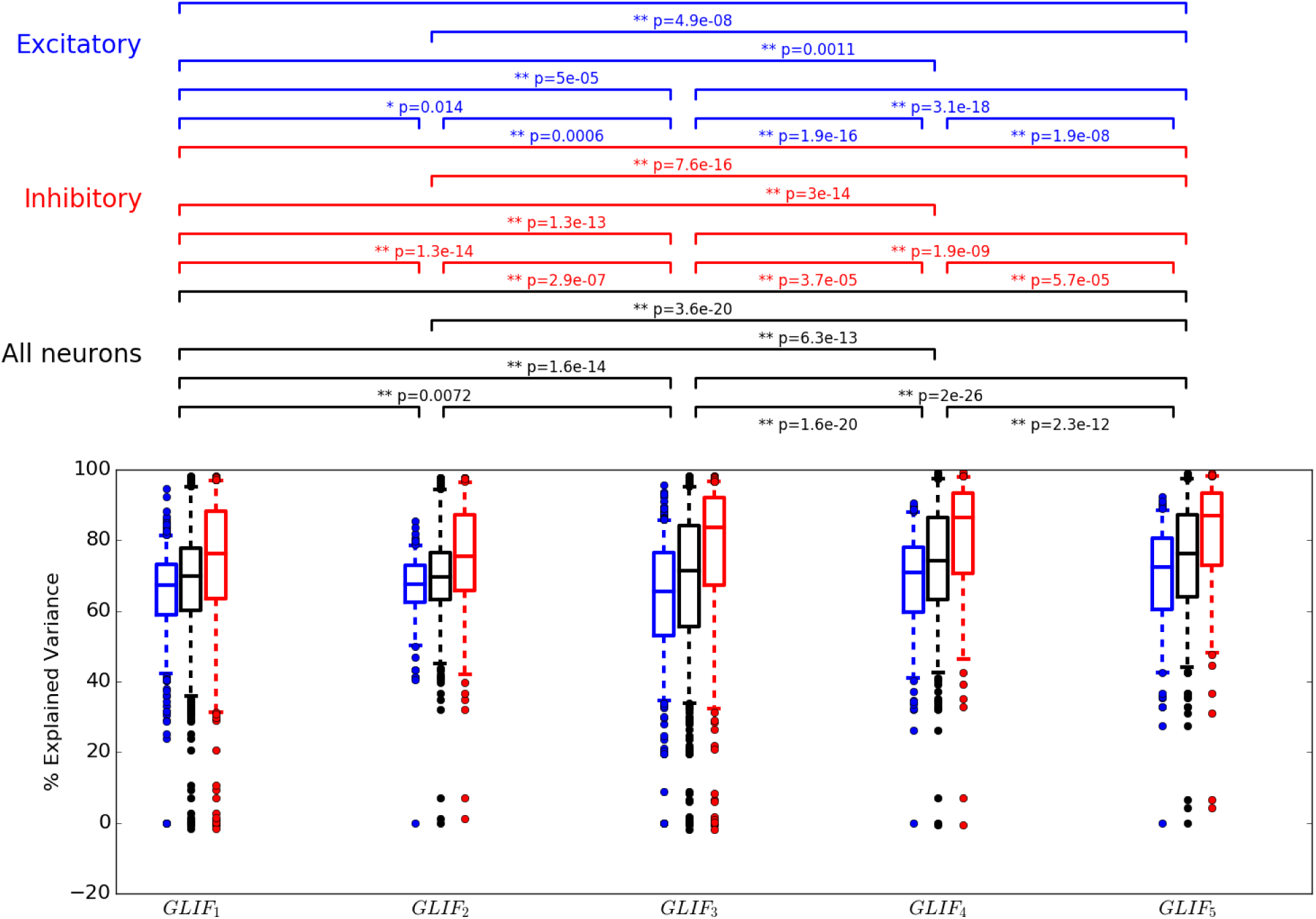
Model performance improves most signi cantly with *GLIF*_4_ model. Explained variance summary for GLIF models for all 771 neurons (black), all 315 inhibitory neurons (red) and all 456 excitatory neuron (blue). Box plots display median, and quartiles. Whiskers reach to 5% and 95% quartiles. Individual data points lie outside whiskers. Brackets above plot denote significant difference between distributions assessed via a Wilcoxon sign ranked test: a single asterisk (*) represents a p-value < 0.05, a double asterisk (**) represents a p-value < 0.01. Distribution values are available in Table 4

### 2.5 Example neurons

In order to provide examples of models that do well reproducing the spike times of the different transgenic cell lines, for each line we select the model which has the highest explained variance from neurons that have all five model levels. Example neurons are labeled with stars in all figures (Figures 7 and 9, and 10 and can be found on the website via the following specimen IDs: Pvalb: 481018305 *GLIF*_4_, Rorb: 314822529 *GLIF*_5_, Ctgf: 510106222 *GLIF*_4_, Vip: 509604672 *GLIF*_5_, Ndnf: 527464013 *GLIF*_5_, Cux2: 490376252 *GLIF*_5_, Chat: 519220630 *GLIF*_4_, Ntsr1: 485938494, *GLIF*_3_, Scnn1a-Tg2: 490205998 *GLIF*_5_, Scnn1a-Tg3: 323834998 *GLIF*_3_, Nr5a1: 469704261 *GLIF*_5_, Htr3a: 477975366 *GLIF*_5_, Sst: 313862134 *GLIF*_5_, Rbp4: 490944352 *GLIF*_5_. Although these neurons are the best performers they may not be the best representational cells. Each of these models have at least one parameter which lies outside the 5 to 95 percentiles for the cre line. To facilitate the choice of other good representative models, neurons that contain all *GLIF* model levels and all model parameters lie within 5 to 95 percentiles are available in Table 5 of the Supplementary Material.

### 2.6 Clustering

In order to assess how well clustering on *GLIF* parameters can potentially classify cell types, we perform a set of clustering experiments using the parameters obtained from the *GLIF* models. We compare the number of clusters from model results with those obtained from clustering the associated features extracted directly from the electrophysiological traces. We assess how well the putative clusters agree with known cell type information based on the transgenic lines from which the cells were derived. To perform clustering a large data set is required. Therefore we use only parameters that are available for all 771 cells. Table 1 describes the parameters used in each clustering paradigm.

**Table 1:** Summary of GLIF models and results. “GLIF” column reports the model level. The “Num. cells” column reports the number of cells for which a GLIF model is constructed. The variables for each model level are listed in the “Variables” column. The parameters fit for each model level are listed in “Model parameters”. Note that resistance was fit along with after-spike currents in models where after-spike currents were implemented. *R*, denotes the resistance fit without ASC and *R_ASC_* denotes the resistance fit along with ASC. Clustering is only performed on parameters that are available for all 771 neurons. Therefore, the’Parameters in clustering’ list is a subset of the total model parameters available for any level. *GLIF*_5_ does not report clusters because there are no additional parameters available for clustering than in *GLIF*_4_. We are unable to cluster on the time scales of every ASC alone as there were five discrete possible values but only two were chosen for each neuron. Therefore, we cluster on the total charge deposited over short *δI_1_/k_1_* and long *δI_2_/k_2_* time scales (continuous numbers) for the model levels that contain after-spike currents. The average explained variance at a time resolution of 10 ms for all neurons at each level is reported as well as the number of clusters that were found using the aforementioned clustering parameters. In the last row, optimized parameters available for all neurons were used in clustering.

| GLIF | Num. cells | Variables | Model parameters | Parameters in clustering | Explained Variance $\Delta t=10$ ms | Num. clusters |
| --- | --- | --- | --- | --- | --- | --- |
| 1 | 771 | $V(t)$ | $R, C, E_l, \Theta_\infty, \delta t$ | $R, C, E_l, \Theta_\infty, \delta t$ | 70.0 | 8 |
| 2 | 289 | $V(t), \Theta_s(t)$ | $R, C, E_l, \Theta_\infty, \delta t, f_v, \delta V, b_s, \delta \Theta_s$ | $R, C, E_l, f_v, \delta V, \Theta_\infty, \delta t$ | 69.7 | 9 |
| 3 | 771 | $V(t), I_1(t), I_2(t)$ | $R_{ASC}, C, E_l, \Theta_\infty, \delta t, k_1, \delta I_1, k_2, \delta I_2$ | $R_{ASC}, C, E_l, \Theta_\infty, \delta t, \delta I_1/k_1, \delta I_2/k_2$ | 71.3 | 8 |
| 4 | 289 | $V(t), \Theta_s(t), I_1(t), I_2(t)$ | $R_{ASC}, C, E_l, \Theta_\infty, \delta t, f_v, \delta V, b_s, \delta \Theta_s, k_1, \delta I_1, k_2, \delta I_2$ | $R_{ASC}, C, E_l, \Theta_\infty, \delta t, f_v, \delta V, \delta I_1/k_1, \delta I_2/k_2$ | 74.3 | 9 |
| 5 | 288 | $V(t), \Theta_s(t), I_1(t), I_2(t), \Theta_v(t)$ | $R_{ASC}, C, E_l, \Theta_\infty, \delta t, f_v, \delta V, b_s, \delta \Theta_s, k_1, \delta I_1, k_2, \delta I_2, a_v, b_v$ | N/A | 76.2 | N/A |
| all param | N/A | N/A | N/A | $R, R_{ASC}, C, E_l, \Theta_\infty$ (from GLIF 1 & 3), $\delta t, f_v, \delta V, \delta I_1/k_1, \delta I_2/k_2$ | N/A | 18 |

We use an iterative binary splitting method using standard hierarchical clustering methods (Please see section “4.6 Supplementary Material”), run over 100 iterations to assess cluster robustness. Using only the parameters which characterize the *GLIF*_1_ model results in eight clusters, with two clusters comprising predominantly inhibitory neurons. Using only parameters from the *GLIF*_3_ model, eight distinct clusters emerge. The addition of reset rules, which are included in the *GLIF*_2_ and *GLIF*_4_ models, results in nine distinguishable putative cell types using the same clustering method (Figures available in Supplimentary Material). However, including the full set of all optimized parameters results in 18 clusters (Figure 6a).

**Figure 6:**
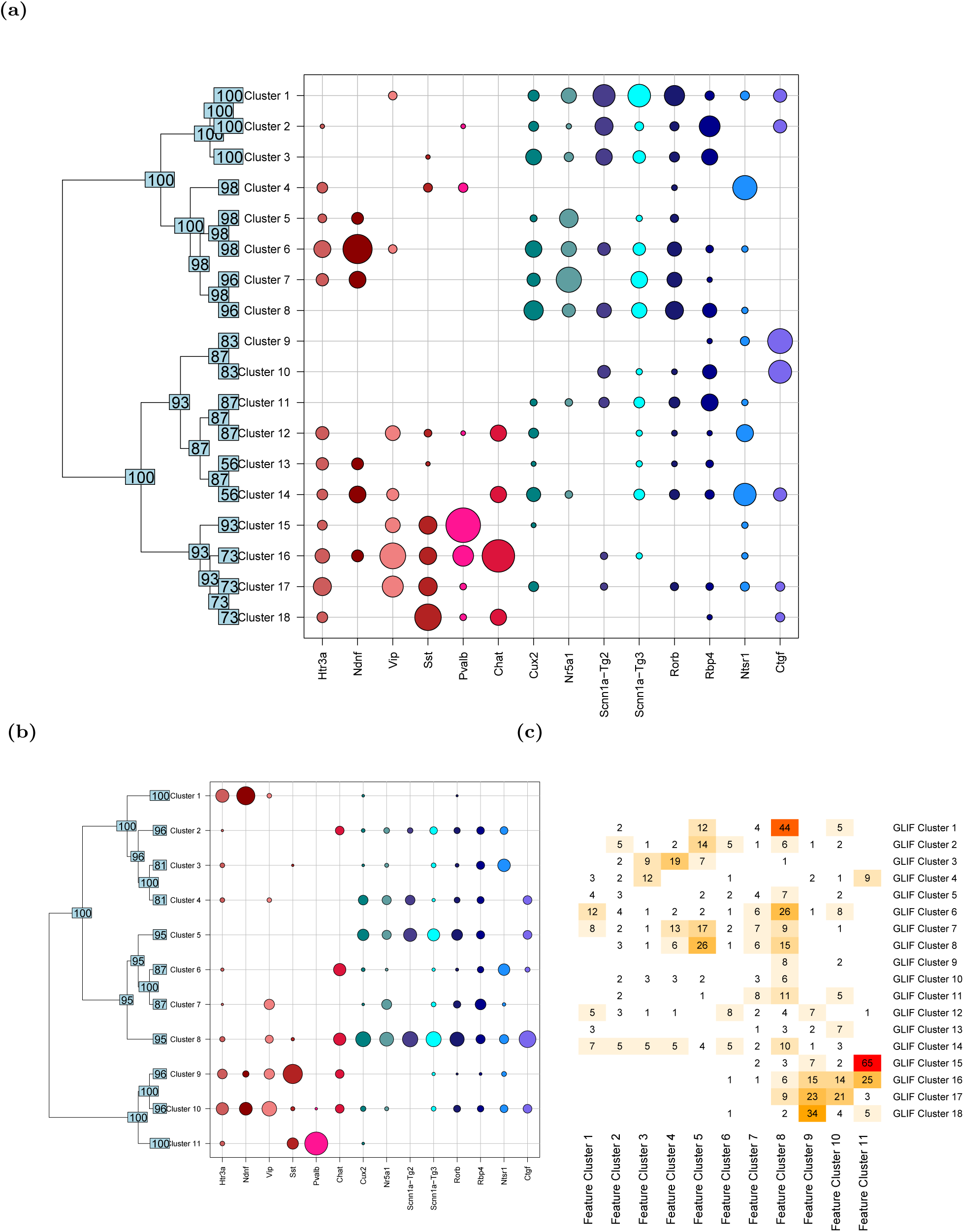
When clustering all 771 GLIF models we identify 18 discrete putative clusters. 11 electrophysiological cell types are obtained clustering based on the features extracted from traces. (a) Summary of clusters obtained by iterative binary hierarchical clustering using all optimized model parameters 8. Each row represents a cluster, and each column a transgenic line. The size of the circle indicates the fraction of cells from a given transgenic line falling into a specific cluster (such that the sum of fractions in a column add up to 1). The dendrogram on the y-axis shows the iterative binary splitting into clusters using the algorithm explained in the text, and the numbers at the leaves and intermediate nodes indicate how often out of 100 clustering runs a given split or terminal cluster was identified. Although the segregation is not perfect, there is separation among lines labeling inhibitory and excitatory cells. In addition, transgenic Cre lines marking Pvalb+, Ntsr1+, and Ctgf+ cells tend to segregate into distinct clusters. (b) A summary of 11 electrophysiological cell types is obtained using the features extracted from traces (Table 10). Each row represents a cluster, and each column a transgenic line. The size of the circle indicates the fraction of cells from a given transgenic line falling into a specific cluster (such that the sum of fractions in a column add up to 1). The dendrogram on the y-axis shows the iterative binary splitting into clusters, as explained in the text, with the numbers at the leaves and intermediate nodes indicating how often out of 100 clustering runs a given split or terminal cluster was identi ed. Although there is separation among lines labeling inhibitory and excitatory cells, the cells from the Ctgf and Ntsr lines are not strongly separated from other cells among the clusters. (c) Schematic of numbers of clusters found with the parameters from each model level. (d) Overlap between GLIF and feature-defined clusters. There is a clear segregation between excitatory and inhibitory neurons. Among excitatory neurons the two classifications are somewhat distinct, with the GLIF clusters coming closer to the laminar distribution of neurons.

The transgenic line composition of each terminal cluster in Figure 6a suggests that clustering based on model fit parameters broadly segregates neurons into previously identified classes of cells. Neurons from transgenic lines labeling predominantly excitatory neurons cluster separately from those labeling mainly interneurons. In addition, sub-categories of neurons appear: these include parvalbumin-positive interneurons, Layer 6a corticothalamic neurons (derived from the Ntsr1 transgenic line), and Layer 6b neurons (derived from the Ctgf transgenic line).

Examining the electrophysiological features of the recorded cells found in the clusters based on the full set of GLIF parameters reveals distinctive characteristics between some of the clusters (Table 10). Cluster 15 contains most of the parvalbumin-positive neurons and accordingly demonstrates many features associated with fast-spiking interneurons: a low action potential upstroke/downstroke ratio, a steep slope of the f-I curve, and a low membrane time constant. Clusters 16, 17, and 18 contains many of the other inhibitory interneurons from the Sst and Vip transgenic lines; these clusters exhibit higher median input resistances, the low upstroke/downstroke ratios (although not as low as the aforementioned cluster 15), and a relatively deep trough following the action potential. Interestingly, many of the cells from the Ndnf transgenic line (which labels inhibitory neurogliaform cells) are found in cluster 6 (and neighboring clusters 5 and 7) on the predominantly excitatory side of the tree. Cluster 6 has the lowest median membrane time constant on that side of the tree, and it exhibits a higher upstroke/downstroke ratio and longer average interspike interval than the other large inhibitory clusters described here.

Among the excitatory-neuron-dominated clusters, there is more similarity among the electrophysiological features, but some relevant differences can be observed. Clusters 9 and 10 (containing many neurons derived from the Ctgf transgenic line) exhibit some of the highest median upstroke/downstroke ratios across all clusters. Cluster 4 (containing many neurons derived from the Ntsr1 transgenic line) has neurons with a shorter membrane time constant, lower upstroke/downstroke ratio, and higher rheobase (I_threshold_ on long steps) than most other excitatory-dominant clusters. Clusters 5, 6, and 7 contain many of the excitatory cells that exhibit bursting behavior. Cluster 3 is notable for its low median input resistance, the shallow median slope of the f-I curve, and relatively high spike-frequency adaptation.

Interestingly, when the same clustering technique is applied directly to the electrophysiological feature data set (Figure 6b), the correspondence between clusters and the excitatory transgenic lines is somewhat weaker. In particular, the layer 6 neurons that emerged in separate clusters with the model parameters are not as well isolated into distinct groups here. Clusters 9, 10, and 11 appear to separate interneurons from the Sst, Htr3a/Vip, and Pvalb transgenic lines relatively well, while cells from the Ndnf transgenic line again cluster more closely with the excitatory cells. Overall, this suggests that dimensionality reduction onto the space of model fit parameters allows for discriminability similar to that obtained by the extraction of a subset of electrophysiological features.

## 3 Discussion

In the adult cortex, the majority of communication between neurons is via chemical synapses from axons onto dendritic or somatic membrane (with a fraction of inhibitory neurons coupled by gap junctions as notable exceptions). The response of these synapses is generally dependent only on the action potentials generated by the presynaptic cell. Thus, we focus on reproducing the temporal properties of spike trains using computationally compact simple GLIF models. We show that a) additional mechanisms included in GLIF models provide and increased explanatory power for test stimuli and b) that the low dimensional description generated by the models capture enough of the complexity of the neuron behavior such that the number of cell types obtained from clustering model parameters is modestly larger than the number of cell types resulting from clustering electrophysiological features.

### 3.1 Model performance of spike time reproducibility

The performance of different models is dependent on the properties of the input currents i.e. if there are large variations in the input current, the capacity to predict the spiking output is better. We chose to compute the performance metrics on stimuli which have the same coefficient of variation 0.2 as the currents measured with slice patching, and a 
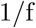
 (pink noise) power distribution of frequencies between 1 and 100 Hz, using two supra-threshold amplitudes. It is important to note that exact comparison of explained variance for models across different publications is difficult, as having stimuli with large temporal variations can lead to more precise firing (Mensi et al., 2012; Pozzorini et al., 2015).

After optimizing the neuronal parameters we were surprised on how well the leaky integrate and fire neuron model performs under such naturalistic conditions, explaining 70.0% of the variance on average. While the reset rules and the afterspike currents separately do not directly improve the model performance, together they lead to a small but signi cant increase to 74.3%. There is a further small but significant improvement, to 76.2%, with the introduction of the voltage-dependent threshold. While the model complexity increases, the fraction of explained variance is computed on a hold-out data-set, not the training data, such that over-fitting on the training data would result in a decreased test explained variance.

Of course, there are limitations to our models: the traces which are reproduced are coming from a current injection into the soma of the neuron in a stereotyped recording condition. We try to keep the statistics of the test stimulus similar to in vivo patch clamp currents, but in vivo additional large sources of variability will occur from stochastic transmission at synapses, complex dendritic integration, neuromodulation etc. These mechanisms, which are necessary for exact integration of our models into systems models are beyond the scope of the current study.

### 3.2 Neuronal Types

In the cerebral cortex there is a wide diversity of neuron types observed at transcriptomic levels Tasic et al. (2016). The existence of different types may be driven by a need to develop particular cell type specific connectivity, neuromodulation, or perform different cell specific computations. When constructing a system level model, it is important to know how many types of neurons are needed to describe the relation between an input current and the resulting output spiking pattern measured in electrophysiological experiments.

In an attempt to characterize cell types associated with the input/output transform measured by electrophysiological experiments, a clustering can be run on a set of electrophysiology features. However, it is not entirely clear which are the most important features to consider. An alternative method is use the entire spike train by synthesizeing the input/output relationship into a model and then performing the clustering on the model parameters. A seemingly intuitive model to use for such a clustering would be biophysically detailed model, as the parameters map well to biological mechanisms. However, a problem with such models is that they do not provide unique solutions: repeated optimization with the same input can lead to solutions which are far away in parameter space. Thus, another approach is to use simpler linear models which have both a) unique parameter solutions, and b) have been shown to reproduce spike times of biological data (Pozzorini et al., 2015). Indeed, previous studies have touched upon the potential for clustering using simplified models (Dong et al. (2013); Mensi et al. (2012)).

Clustering upon 11 fit and optimized GLIF parameters available for all 771 neurons (Table 1) we obtained 18 clusters using a simple iterative hierarchical clustering method. We were surprised at this large number considering that we started clustering with cells from 14 transgenic lines (all transgenic lines for which we had less than 7 neuron models were excluded) and only 11 clusters were obtain using 16 electrophysiological parameters (Supplementary Material Table 10). This suggests that the GLIF model parameters have at least a similar capacity of separating distinct clusters of cell as the features especially designed to do that. The unsupervised clusters do not perfectly match the transgenic lines, which is not surprising given that the transgenic lines are known to comprise multiple molecularly defined cell types. However, the segregation of neurons into putative types based on the model fit parameters does recapitulate some expected differences among classes of neurons based on other data modalities.

It seems that the mechanism which is most responsible for the increase in the capacity to classify cells is combination of the resistances fit both with and without afterspike currents and the optimized *th*_∞_ for *GLIF*_1_ and *GLIF*_3_ models (optimized *th_∞_* values for *GLIF*_2_, *GLIF*_4_, and *GLIF*_5_ are not available for all neurons).

### 3.3 Conclusion

Here we conclude that GLIFs are an exceptionally useful tool for single neuron models. They can simultaneously reproduce the spike times of biological neurons and reduce the complex mapping between the input current to the spike train output to a small number of parameters. These parameters synthesize the richness of the observed neuronal behaviors to allow the same number of classes to be obtained from models as there are from features characterizing the data.

We provide a database of GLIF models that can facilitate the development of system models with cell types available on the Allen Cell Types Database at http://celltypes.brain-map.org/.

## 4 Supplementary Material

### 4.1 Model Definitions

The five different GLIF models are described below, in order of increasing level of complexity, with their evolution equations and reset rules. For all the models, *C* represents the membrane capacitance, *R*, is the membrane resistance, *E_L_* is the resting potential and *I*_*e*_(*t*) is an external current injected into the cell. *t*_+_ and *t*_-_ represent the time just after and before a spike respectively.

#### 4.1.1 Leaky Integrate-And-Fire (*GLIF*_1_)

The traditional Leaky Integrate-And-Fire (LIF) neuron is a hybrid system characterized by an evolution equation for the membrane potential V(*t*). We refer to this model as *GLIF*_1_ as it is the starting point for further GLIF models.

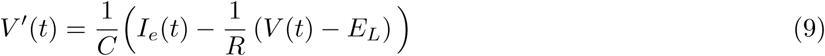

When the membrane potential becomes larger than a threshold *V*(*t*) > Θ (which in this case is just the instantaneous threshold, Θ_∞_), a reset rule is evoked which sets the membrane voltage to a reset voltage *V_r_*.

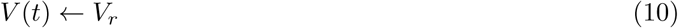

#### 4.1.2 Leaky Integrate-And-Fire with biologically de ned reset rules (*GLIF*_2_)

*GLIF*_2_ has an evolution equation for two state variables, the membrane potential *V*(*t*) and a spike-dependent threshold component Θ_s_(*t*) which is updated by spikes and decays back to zero with a time constant 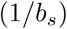

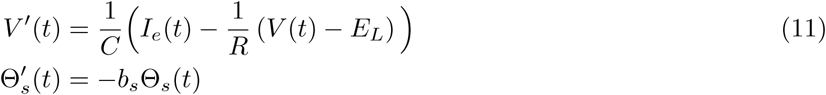

along with a reset rule for both voltage and the spike-dependent threshold if the membrane potential becomes larger than a threshold 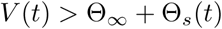

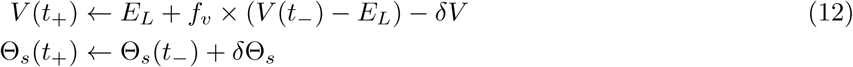

The update to the spike-dependent component of the threshold is additive, with *δ*Θ_*s*_ added after every spike. The update to the membrane potential has a multiplicative coeffcient *f_v_* and and an additive constant *δV*

#### 4.1.3 Leaky Integrate-And-Fire with after-spike currents (*GLIF*_3_)

In GLIF models, the rapid membrane potential fluctuation due to the fast voltage-activated (i.e. sodium and potassium) ion currents during a spike, is considered separately from the slow, subthreshold region of the membrane potential and is incorporated into the reset rules. However, ion currents activated by a spike could have effects over longer time scales. After the stereotypical, sharp membrane potential transitions during an action potential are eliminated, the longer term effects are modeled as additional currents *I_j_* with pre-de ned time constants 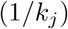,

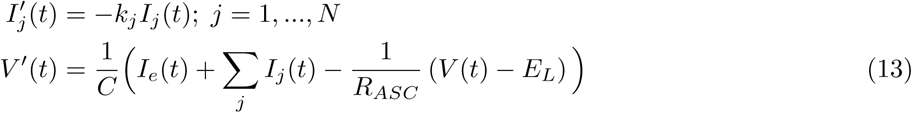

Here *R_ASC_* is used to explicitly point out that the resistance is fit along with the after spike currents. The update rule, which applies if *V*(*t*) > Θ_∞_, is given by:

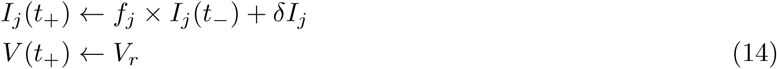

with the currents updated using a multiplicative constant 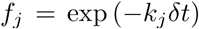(because after spike currents decay thought the spike cut length) and an additive constant, *δI_j_*.

#### 4.1.4 Leaky Integrate-And-Fire with biologically de ned reset rules and after-spike currents (*GLIF*_4_)

Combining the biological reset rules model, *GLIF*_2_ with the after-spike current model *GLIF*_3_ described above gives a model defined by:

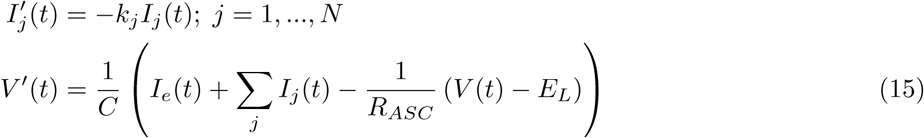

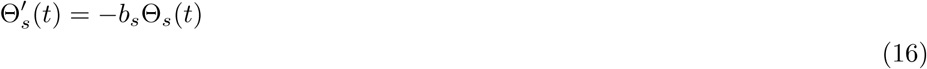

The update rule, which applies if 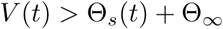,

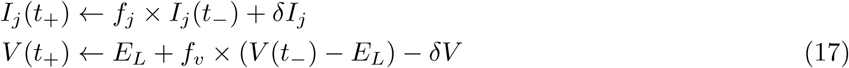

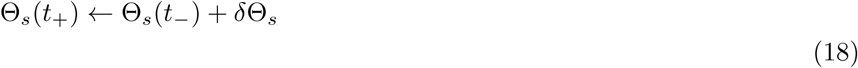

#### 4.1.5 Leaky Integrate-And-Fire with biologically de ned reset rules, after-spike currents and a voltage dependent threshold (*GLIF*_5_)

For the *GLIF*_5_ model, the state variables are the membrane potential *V*(*t*) (as in all *GLIF* models), a threshold component which is evoked when there is a spike (Θ*_s_*) (as introduced in *GLIF*_2_), a set of after-spike currents *I_j_*) (as introduced in *GLIF*_3_), and an new additional threshold parameter that is dependent of the membrane potential (Θ*_v_*). It is assumed that these state variables evolve in a linear manner between spikes:

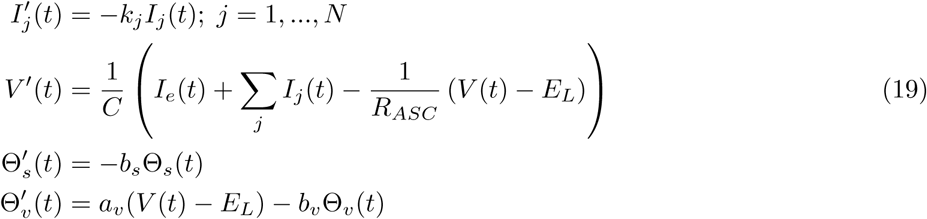

Where 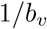 is the time constant of the voltage-dependent component of the threshold and, *a* can be interpreted as a’leak-conductance’ for the voltage-dependent component of the threshold.

If 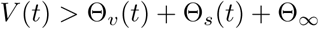a spike is generated and the state variables are updated:

**Table 2:** Variable description

| Mechanism | Symbol | Variable |
| --- | --- | --- |
| | $V(t)$ | Membrane Potential |
| ASC | $I_j(t)$ | After-spike currents |
| reset | $\Theta_s(t)$ | Spike-dependent thresh-<br>old component |
| adapt. th. | $\Theta_v(t)$ | Voltage-dependent<br>threshold component |

**Table 3:** Parameter description

| Mechanism | Symbol | Parameter | Fit From | Post-hoc<br>Optimized |
| --- | --- | --- | --- | --- |
|  | C | capacitance | V during low amplitude noise | No |
|  | G | membrane conductance | subthreshold V during noise | No |
| | $E_L$ | resting potential | resting V before noise | No |
| | $\Theta_\infty$ | instantaneous threshold | short square input | Yes |
| reset | $f_v$ | voltage fraction following reset | all noise spikes | No |
| reset | $\delta V$ | voltage addition following reset | all noise spikes | No |
| reset | $b_s$ | spike-induced threshold time constant | triple short square | No |
| reset | $\delta\Theta$ | threshold addition following reset | triple short square | No |
| ASC | $\delta I_j$ | afterspike current amplitudes | subthreshold V during noise | No |
| ASC | $k_j$ | afterspike current time constants | subthreshold V during noise | No |
| adapt. th. | $a_v$ | adaptation index of threshold | prespike V during noise | No |
| adapt. th. | $b_v$ | voltage-induced threshold time constant | prespike V during noise | No |

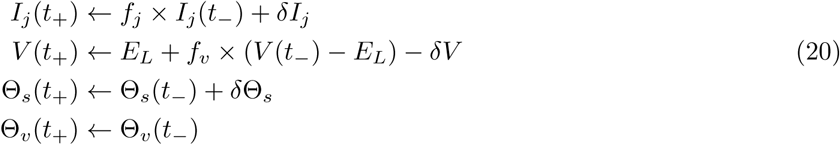

For all state variables *X_k_*(*t*), the value immediately after the spike, *X_k_*(*t*_+_) was related to the value immediately before the spike, *X*_*k*_(*t*) via a set of update parameters: *f_k_* represented the fraction of the prespike value of *X_k_* which is maintained after the spike and δ*X_k_* represents the values updated by a spike.

### 4.2 Parametertting and distributions

Model parameters are extracted from the electrophysiology data as described below. Python code for all algorithms can be found in the Allen Software Development Kit (SDK) on the Allen Cell Types Database website. All code was executed using Python 2.7. In Figures 7, 8 and 9, and 10 we show slices of the parameter space obtained via fitting.

**Figure 7:**
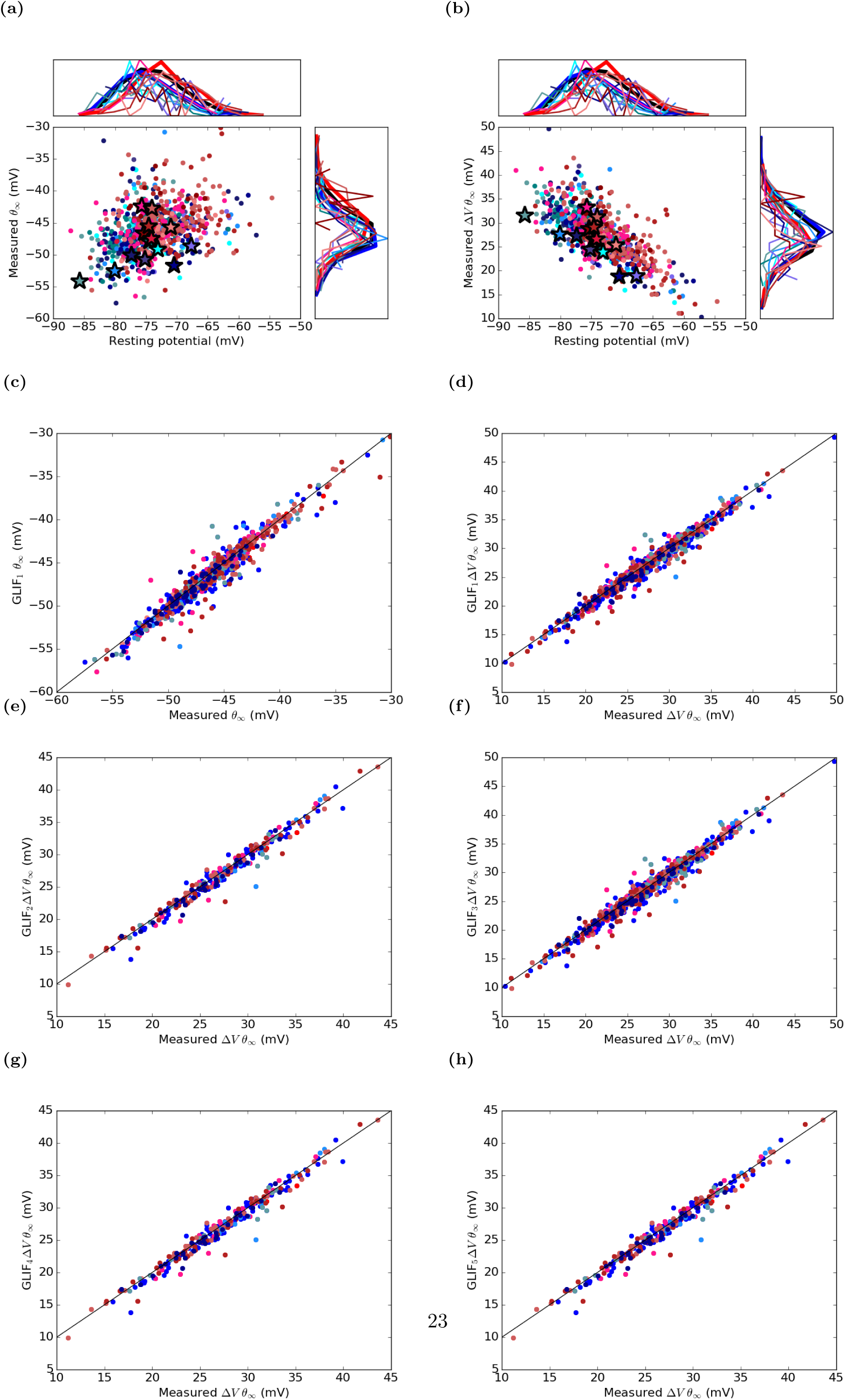
Resting potential and instantaneous threshold for biologically measured values and post-hoc optimized models. In (a) absolute threshold is shown, in (b) threshold is in reference to the resting membrane potential for the biologically measured threshold from the short square stimulus (Figure 2 Stimuli). This gure is also shown in the main text (Figure 3) (c) and (d) show the similarity between threshold values before and after post-hoc optimization for absolute threshold and (c) and relative threshold (d). (e), (f), (g), and (h) compare the biologically measured relative threshold versus the relative threshold obtained by optimizing _1_ of (e) *GLIF*_2_, (f) *GLIF*_3_, (g), *GLIF*_4_, and (h) *GLIF*_5_. Black lines represents unity. Shades of blue represent excitatory, transgenic lines, shades of red represent inhibitory lines. The full list of colors corresponding to specific transgenic lines can be found in Table 2 Transgenic Lines. Normalized histograms of the data are shown in the side panels. The thick black histogram represents data from all transgenic lines, the thick blue line represents all data from excitatory and the thick red represents inhibitory lines. Stars denote example neurons shown throughout the manuscript. All neurons with a measured instantaneous threshold below −60 mV were globally excluded from the data set.

*Spike initiation time and threshold [t_s_, Θ_bio_]:* Spikes are typically detected by the following procedure. The upstroke of a spike is defined as the portion of the action potential wave form between the instant at which the rise in potential for a given time exceeds 20 mV/ms and the peak of the action potential. For each neuron the average value of the maximum dV/dt during the upstrokes of all action potentials is calculated. The time of spike initiation and threshold is then defined by the time and voltage at which dV/dt reaches5% of the average maximum dV/dt during an upstroke.

For stimuli that involve a short, intense current pulse stimulation, the passive rise in voltage preceding a spike can exceed the standard 20 mV/ms initial detection value, leading to inaccurate spike time identification. In those cases, the maximum dV/dt is measured during the current pulse, and the spike detection value is adjusted to be 10% higher than that. In addition, the threshold identification value (typically 5% of the average maximum dV/dt) is also adjusted to ensure that it is at least 20% higher than the dV/dt at the end of the stimulus pulse.

*Spike cut length and voltage reset [*δt*, *V*_*t*+_]:* GLIF models aim to reproduce the timing of the spikes via sub-threshold data (not the voltage waveform dictated by the highly non-linear ion fluctuations during the action potential). Therefore, the action potential waveforms are removed from the voltage trace. Here we use a principled way to estimate the duration of the spike that is removed and the reset voltage in models where biological reset rules are implemented (*GLIF*_2_, *GLIF*_4_, and *GLIF*_5_). Action potentials in a neuron have stereotyped shapes with a sharp rise, followed by a hyperpolarized refractory period and a return to a voltage that is most often lower than the voltage before the spike (voltage reset). As shown in Figure 8, we align all of the action potentials of the training noise data (noise 1) to the spike initiation and ask at what time within a window of 1 to 10 ms after spike initiation does the post spike voltage best predict the voltage at spike initiation given a linear dependence. i.e. a line is fit between the voltages at spike initiation (prespike voltages) and the voltages at each time point after spike initiation (postspike voltages). The fit which minimizes the residuals is chosen to define the time at which the spike ends. In all models, the duration of the spike is removed from the voltage traces. For the *GLIF*_2_, *GLIF*_4_, and *GLIF*_5_ models, the postspike voltage is reset via the best t linear model 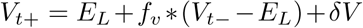, where V_t−_ is the voltage of the model before the spike, *δV* is the voltage intercept, *f_v_* is the slope of the fit, and *V_t+_* is the voltage of the model after the spike.

**Figure 8:**
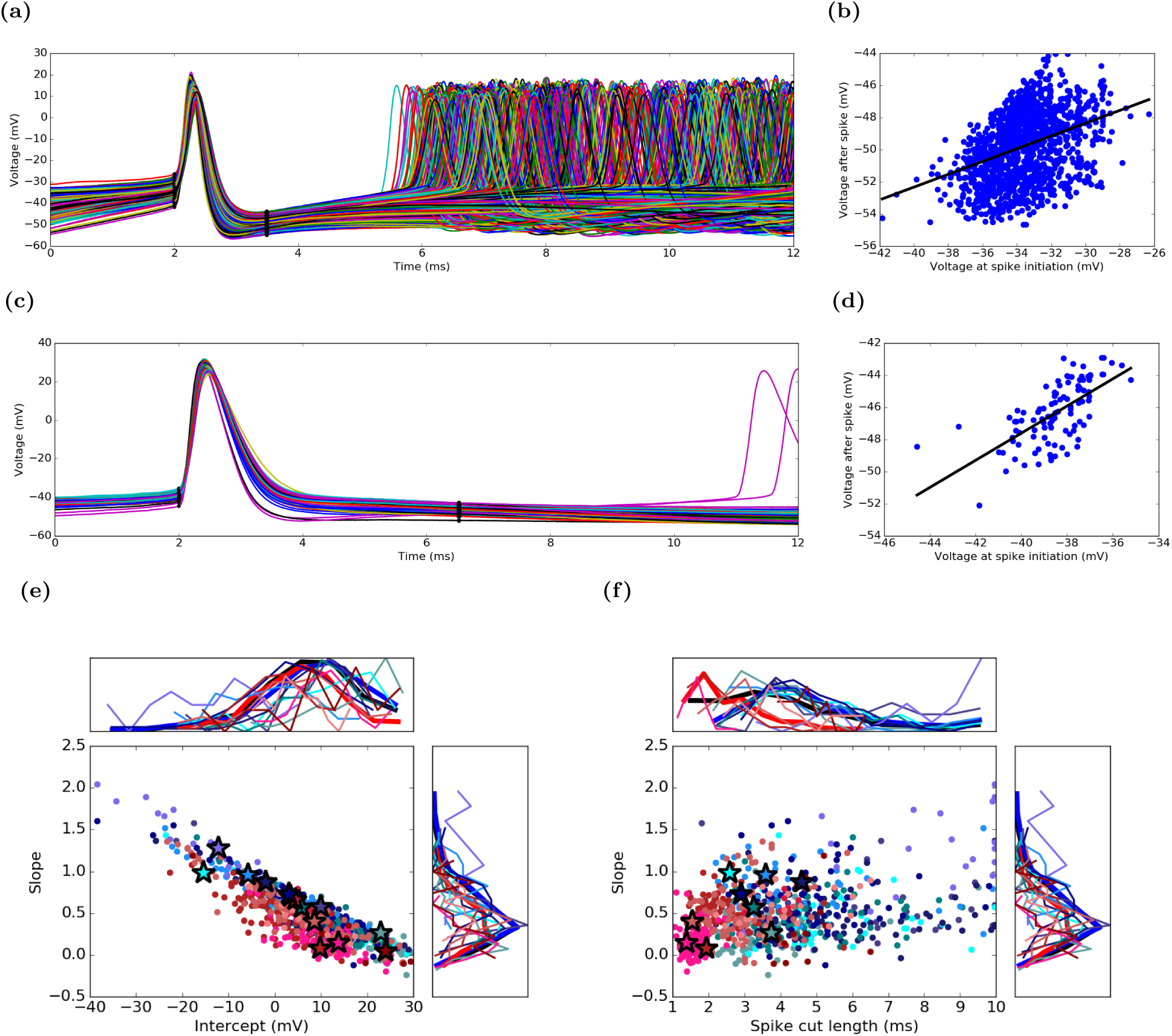
The action potentials waveforms are excluded from model fitting but are used to derive voltage reset rules. (a-d) illustrate examples of the spike cutting and reset rule calculation from two example neurons (a, b are from the Htr3 477975366 neuron; c, d are from the Rorb 314822529 neuron) from two di erent lines. (a, c) All spikes from the noise 1 stimuli (Figure 2 Stimuli) are aligned to spike initiation. Here di erent colors represent di erent spikes of one neuron (not di erent transgenic lines). Dots represent spike initiation and termination found by minimizing the residuals from a linear regression between pre-and post-spike voltage in a window 1 to 10 ms after spike initiation (b, d). In the (*GLIF*_2_, *GLIF*_4_, and *GLIF*_5_ models, voltage reset is calculated by inserting the model voltage when it reaches threshold into the equation de ned by the line (Equations 5, 12, 17 and 20). (e) and (f) show distributions for slope, intercepts, and the spike cut length for all neurons. Normalized histograms of the data are shown in the side panels. All neurons with an intercept < 30 mV were globally excluded from the data set. (f) is a replicate of the figure shown in the main article (Figure 3).

*Instantaneous threshold [Θ_∞_]:* All models (neurons) spikes when the voltage crosses threshold. An estimate of the instantaneous threshold, Θ_∞_, is the voltage at spike initiation of the lowest amplitude supra-threshold short square (Figure 2 Stimuli). Threshold is a critical parameter a ecting the general excitability of the cell. The relationship between voltage and stimulus likely becomes non-linear near threshold. Because small errors in Θ_∞_ measurement will greatly influence spiking behavior Θ_∞_ is tuned during the post-hoc optimization of every model such that the likelihood of the observed spike train in the training stimulus is maximized.

*Resting potential [E_l_]:* The resting potential was defined as the mean resting potential of the noise 1 sweeps calculated by averaging the pre-stimulus membrane potential.

*Resistance [R], Capacitance[C], and afterspike currents [I_j_(t)]:* Because the GLIF model is linear, many of the parameters can be fit via linear regression on subthreshold data similar to the methods in Pozzorini et al. (2015). Initially, R and C are calculated via a linear regression of the subthreshold noise in the first epoch of noise stimulation within the three-epoch noise sweeps (this stimulus does not elicit any spikes) as observed in Figure 9d. The neurons which elicited spikes during the subthreshold stimulation were eliminated from subsequent fits. Please see Supplementary Material for example calculations. These values of resistance are used for models that do not include after-spike currents (*GLIF*_1_, *GLIF*_2_).

For the GLIF model levels where after-spike currents are included (*GLIF*_3_, *GLIF*_4_, *GLIF*_5_), after spike currents and resistance are fit via a linear regression on the periods between the spikes in the supra-threshold noise data. During this calculation, the capacitance and resting potential are forced to the previously calculated values. We choose to force capacitance while continuing to fit resistance because capacitance is a consistent property of the membrane which does not change based on the state of a neuron. However, the resistance of membrane will depend on the state of a neuron and will most likely differ with the activation of ion currents whose subthreshold affects are models here with after-spike currents.

After spike currents are modeled using exponential decaying basis functions with an amplitude and a time scale (the dynamics of the exponential functions are described in equation 15). The time scales and amplitudes are obtained by providing two from a set of five basis currents with varied time scales 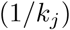 ([3.33, 10, 33.3, 100, 333.33] ms) to a generalized linear model (GLM). The GLM (implemented via Python’s statsmodels.api. GLM) is used to calculate the amplitudes corresponding to each basis current and the resistance by regressing the sum of the derivative of the voltage and the external current divided by the capacitance 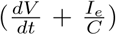 against the basis currents and the leak term. *C* and *E_L_* are calculated as mentioned in the above text. The GLM is run with all combinations of two of the five possible time scales leading to a choice among 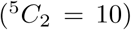 pairs of basis currents. The optimal pair of basis currents that yield the maximum log-likelihood value are chosen. The total charge, *Q*, deposited by one after-spike current in the neuron per spike is 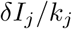. Since total charge is a biologically relevant parameter and it is easier to perform clustering on continuous values, we often use the total charge instead of the individual parameters, *δ_I_*, and, *k_j_* (which is discrete), represented in the equations (Figures 3 and 9, and Tables 6 and 8)

*Spike initiated component of the threshold [ Θ_s_]:* The evolution of the threshold of (*GLIF*_2_, *GLIF*_4_, and *GLIF*_5_ contains a spiking component of the threshold, Θ_*s*_. This contribution to the threshold represents the effect spiking has on the threshold of the neuron due to the inactivation of a voltage-dependent sodium current. This inactivation can be interpreted as a rise in the threshold of the neuron, and the movement from the inactive to a closed state can be modeled as a linear dynamical process. The change in threshold is fit with an exponential, and the values of its amplitude and time constant are calculated from the triple short square sweep set data as seen in Figure 10. For each pulse in the triple short square data that produces a spike, the voltage at spike initiation (the threshold) is calculated. The mean of the threshold of the first spike of each individual triple short square stimulus was taken to be the reference threshold. For all spikes that are not first spikes in each individual triple stimulus, the time since the last spike is calculated (referred to as an ISI) along with the threshold relative to the reference threshold. An exponential is then fit to the ISI versus threshold data. The exponential is forced to decay to the reference threshold.

*Subthreshold voltage dependent component of the threshold [Θ_*v*_]: GLIF*_5_ contains both a spiking component of the threshold, Θ_*s*_, (see above) and a subthreshold voltage component of the threshold, Θ_*v*_. The subthreshold voltage component represents the effect of subthreshold membrane potential on the threshold (e.g., voltage dependence of sodium current reactivation). The voltage component of the threshold evolves according to Θ^’^_*v*_(t) in equation 19. In order to fit *a_v_*, and *b_v_* the sum of the squared error between the voltage component of the _threshold_ of the biological neuron Θ_*vbio*_ and the voltage component of the threshold of the model Θ_*v*_ is minimized. Θ_*vbio*_ is found by subtracting the spike component of the threshold, Θ_*s*_ from the actual value of the voltage at the time of spike initiation Θ_*bio*_. Θ_*vbio*_ = Θ_*bio* – Θ*s*_. Although one could simulate the voltage component of the threshold using Euler’s method, the analytical solution (for derivation please see Supplementary Material) was used to speed up the minimization process. Minimization was preformed using a Nelder-Mead simplex algorithm (fmin in the scipy optimization tool box: scipy.optimize.fmin)).

### 4.3 Evaluation of Model Spike Times

After all parameters of the model were optimized, the time-evolution of the models was determined using an Euler exact method. The spike times of the model were evaluated against the spike times of the neuron using the’explained temporal variance’ metric described below (Figure 4). This metric describes how well the temporal dynamic of the spiking response is captured by the model for a particular level of temporal granularity Δ*t*.

A spike train is represented as a time series, at resolution Δ*t*, of binary numbers. All numbers in a spike train are zero unless a spike occurs: a spike was denoted with a one. Any spike train can be converted into a single peristimulus time histogram, stPSTH, by convolving a spike train with a Gaussian.

In cases with many repeats of the same stimulus, a peristimulus time histogram (PSTH) could be calculated by taking the mean of the stPSTH at each instant in time. If there were n stPSTHi denoted with index i, where i goes from 1 to n, i=1,2…n, the PSTH was equal to the column mean of the stPSTHi:

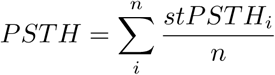

The variance in spiking output of neurons is described by the variance of the PSTH. The explained variance (EV) between any two PSTHs (multiple or single train) is:

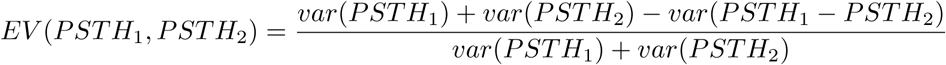

The explained temporal variance by the mean across-trials PSTH of the neuron, *EV_D_* is the upper limit on how well the model can perform:

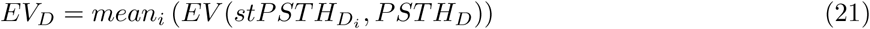

where i denotes the ith data stPSTH and n denotes the total number of data sweeps.

Since the models here were deterministic, the pairwise explained variance of the data with the model is calculated by taking the mean of the explained variance between every data stPSTHD and the model PSTHM.

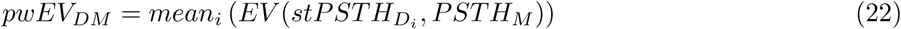

Where i denotes the ith data stPSTH and n denotes the total number of data sweeps.

The ratio of the pwEV of the model to the data versus the pwEV within the data is bounded from below, by 0, where nothing is explained, and from above by 1, explaining the data as best as possible. Explained variance illustrated in Figure 4 for the two exemplary canonical neurons and reported. The ratio of the pwEV of the model to the data versus the pwEV within data at a resolution of Δ_*t*_ = 10 ms converted to a percentage is used as the explained variance metric used throughout the manuscript (Figures 5, 13 and 14 and Table 5)

### 4.4 Post-Hoc Optimization

An additional non-linear optimization step was performed to optimize the instantaneous threshold using a Nelder-Mead simplex algorithm (Python scipy module scipy.optimize.fmin). The instantaneous threshold is an essential parameter governing the excitability of the neuron. As the current/voltage dependence for the neuron becomes highly nonlinear near the threshold, and we rely on linear models, to better fit the excitability of the models we search for the value of the instantaneous threshold which best predicts the spike train during the test stimulus. Comparing Figure 5 of the main text with Figure 11 illustrates the importance of the post-hoc optimization on explained variance.

#### 4.4.1 Forced-spike Model Paradigm

During optimization, each spike should be fit without erroneous after-spike current history affecting the fitting of the subsequent spikes. The model voltage, threshold, and after-spike amplitudes are forced to reset at the time the biological neuron spikes. The reset rules are denoted in the GLIF model section above. When a model is run in its normal forward-running manner, the prespike values of the voltage, *V*(*t*_–_), threshold, Θ(*t*_–_), and AS currents, *I_j_(*t*_–_)*, are the values of the model at the point where the model voltage crosses the model threshold. In the forced spike paradigm, to ensure that the voltage model is reset below the threshold of the model, the prespike voltage was set equal to the prespike threshold, *V*(*t*–) = Θ(*t*_–_) at the time of the neuron spike. Note that this reset only affects *GLIF*_2_, *GLIF*_4_, *GLIF*_5_, where the prespike voltage affects the postspike voltage.

#### 4.4.2 Objective Function: Maximum Likelihood Based on Internal Noise (MLIN)

Similar to a series of previous studies Paninski et al. (2004); Dong et al. (2011); Mensi et al. (2012); Pozzorini et al. (2015), the likelihood that the observed spike train was obtained by the model was maximized. However, the exact method of constructing the likelihood was different in that the noise is not tuned but rather uses a direct estimate of the biological neurons internal noise (Maximum Likelihood based on Internal Noise) was used.

Because these GLIF models are deterministic, estimating the likelihood required adding a source of noise external to the model. Rather than searching for the parameters of this noise, a parametric description of the internal noise of each neuron is used (Figure 12). As this internal noise could depend on the membrane potential, and the most relevant potential is near threshold, the variation in membrane potential during the steady state period of the largest subthreshold square pulse response is characterized.

The probability density of the neuron being at a potential v away from its mean is fit with an exponentially decaying function:

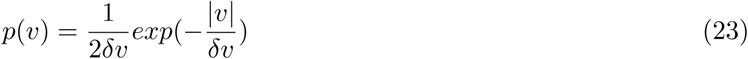

This variation was considered to be additive to the membrane potential of the deterministic neuron. This allowed the computation of the probability that a neuron with the given noise would produce a spike if the deterministic model had a difference between threshold and membrane potential:

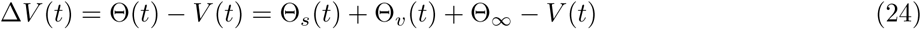

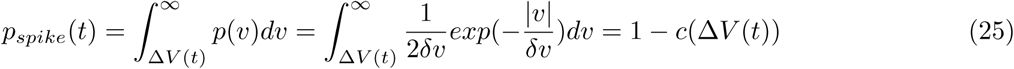

where *c* represents the cumulative distribution of the intrinsic noise.

The likelihood of the model producing a set of spikes at the times the biological neuron produces them is:

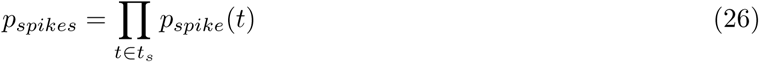

However, the model neuron must both produce spikes when the biological neuron does, and not produce spikes when the biological neuron does not.

The likelihood of a model neuron not producing spikes is not independent for two nearby time points, as the intrinsic noise has a nontrivial autocorrelation. To estimate the likelihood of the model neuron not producing spikes at the times the biological neuron does not produce spikes, one sample for the internal noise is drawn for each time period of the autocorrelation, and following this time scale a new independent sample is drawn. As such, the inter-spike intervals are binned with bin sizes equal to the autocorrelation (typical autocorrelations time scales are much smaller than the inter-spike intervals). The grid times start from a spike time and advance by the autocorrelation time scale, ending at a predefined short time (5 ms) before the next spike. To estimate the likelihood of a neuron not producing a spike within a bin, the minimal difference between threshold and membrane potential within the bin is chosen which generates the grid differences:

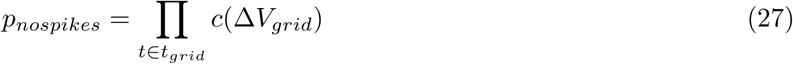

The log likelihood of the entire spike train being exactly reproduces was

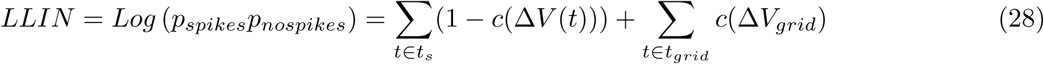

#### 4.4.3 Simplex Method

The instantaneous threshold, Θ_∞_ is optimized to minimize MLIN using a simplex algorithm (Nelder-Mead). During this procedure Θ_∞_ is itself is not optimized, instead, a multiplicative factor (coefficient) is optimized. To ensure that the optimization routine did not return a sub-optimal local minimum, the overall optimization is rerun three times. Each time the overall optimization is rerun, the coefficients are randomly perturbed between an interval of ±0.3 of their last found optimized value. Within each of the overall reruns, the stability of the convergence is confirmed by reinitializing the algorithm (i.e. re-inflating the simplex) three times at the optimal position in parameter space with a small random perturbation within an interval of ±0.01 and then re-running the simplex.

Because the voltage dynamics are governed by a first-order linear differential equation, the voltage dynamics are forward simulated over a single time step using an exact Euler time stepping method. The time step is chosen to be 0.2 ms. The current that forces the dynamics is averaged across the time step.

### 4.5 Performance of individual transgenic lines

The explained variance of individual transgenic lines are shown in Figures 14 and 13 and 14.

### 4.6 Clustering

In order to cluster cells by model parameters or extracted electrophysiological features, we used an iterative binary splitting method using standard hierarchical clustering methods, as follows:

1. Select parameters to cluster on (See Table 1 in the main text).

**Table 4:** Explained variance of all model levels after post-hoc optimization. The single number in each cell is the median. The low and high quartiles are in brackets below the median. All Cre lines with ten or more neurons (n) present in the LIF level are included. Some neurons have the required stimuli for LIF models but do not have the stimuli required for higher level models. When there are not more than 7 neurons in the level, the values are denoted with a NaN. Data illustrated in Figures(5, 13, and 14)

| | n | $GLIF_1$ | $GLIF_2$ | $GLIF_3$ | $GLIF_4$ | $GLIF_5$ |
| --- | --- | --- | --- | --- | --- | --- |
|  |  | 70 | 69.7 | 71.3 | 74.3 | 76.2 |
| all neurons | 771 | [35.7, 95.3] | [44.4, 94.6] | [33.8, 95.3] | [41.6, 97.5] | [43.3, 97.4] |
|  |  | 76.4 | 75.5 | 83.7 | 86.4 | 87 |
| inhibitory | 315 | [31.4, 97] | [41.6, 96.4] | [31.9, 96.7] | [45.5, 98.1] | [48, 98.2] |
|  |  | 67.3 | 67.7 | 65.5 | 70.8 | 72.6 |
| excitatory | 456 | [42.1, 81.5] | [50.1, 78.7] | [34.4, 85.8] | [40.7, 88.1] | [42.6, 88.8] |
|  |  | 60.4 |  | 53 |  |  |
| Scnn1a-Tg2 | 34 | [45.3, 77.7] | NaN | [33.7, 74.8] | NaN | NaN |
|  |  | 67.7 | 69.5 | 68.7 | 71.6 | 73.2 |
| Nr5a1 | 64 | [48.8, 81.4] | [62.5, 77.3] | [47.3, 82.9] | [55.5, 88.1] | [60.9, 88.7] |
|  |  | 65.6 | 66.3 | 58.3 | 64.6 | 64.6 |
| Scnn1a-Tg3 | 47 | [45.1, 76.5] | [53.9, 80.6] | [36.5, 84.7] | [42.6, 88.3] | [36.5, 87.6] |
|  |  | 65 | 69.2 | 63.4 | 68.1 | 71 |
| Rorb | 108 | [40.8, 81] | [51.1, 78.6] | [21.6, 84] | [38, 84.3] | [43.1, 86.4] |
|  |  | 69.9 | 67.8 | 70.2 | 76.1 | 76.9 |
| Cux2 | 74 | [46.9, 81] | [58.7, 74.6] | [37.1, 86.6] | [50.8, 87.6] | [54, 88] |
|  |  | 71.3 | 65.1 | 79.2 | 79.5 | 81.6 |
| Ntsr1 | 44 | [50.3, 82.3] | [42.8, 81.2] | [42.6, 88.7] | [53.6, 90] | [53.3, 90.7] |
|  |  | 61.2 |  | 48.2 |  |  |
| Ctgf | 21 | [28.9, 73.6] | NaN | [23.5, 75.2] | NaN | NaN |
|  |  | 69.6 | 68 | 66 | 72.7 | 73.8 |
| Rbp4 | 64 | [46.4, 83.9] | [55.6, 78.5] | [34.9, 84] | [47.5, 85.8] | [49.6, 89.2] |
|  |  | 75 | 75.5 | 75.3 | 79.4 | 80.1 |
| Sst | 92 | [19.1, 95.1] | [39.2, 93] | [15.2, 95] | [45.2, 95.9] | [48, 96] |
|  |  | 95 | 94.9 | 94.4 | 97.9 | 97.9 |
| Pvalb | 82 | [29.9, 97.8] | [33.9, 97.8] | [29.5, 97.6] | [38.5, 98.8] | [40.6, 98.8] |
|  |  | 73.2 | 69 | 84.8 | 86.8 | 87.1 |
| Htr3a | 96 | [35.4, 86.1] | [47.6, 87.3] | [41.9, 93.4] | [47.5, 95.1] | [53.6, 95.3] |
|  |  | 80.5 |  | 85.3 |  |  |
| Ndnf | 13 | [67.3, 89.2] | NaN | [69.2, 90.3] | NaN | NaN |
| Chat | 7 | NaN | NaN | NaN | NaN | NaN |
|  |  | 67.5 |  | 72 |  |  |
| Vip | 25 | [34.6, 86.8] | NaN | [36.2, 89.7] | NaN | NaN |

**Table 5:** Neurons which have all 5 GLIF models and all model parameters are within 5 and 95 percentiles within each transgenic line. Columns 1 though 5 denote the explained variance of models 1 through 5. The best explained variance within the set is emboldened.

| Specimen ID | 1 | 2 | 3 | 4 | 5 |
| --- | --- | --- | --- | --- | --- |
| Pvalb |  |  |  |  |  |
| 475545311 | 0.943 | 0.928 | 0.935 | 0.961 | 0.965 |
| 515305346 | 0.945 | 0.978 | 0.944 | 0.982 | **0.984** |
| 481127173 | 0.974 | 0.963 | 0.960 | 0.979 | 0.980 |
| 484742372 | 0.971 | 0.946 | 0.967 | 0.983 | 0.982 |
| 478949560 | 0.922 | 0.940 | 0.927 | 0.942 | 0.943 |
| Rorb |  |  |  |  |  |
| 473020156 | 0.604 | 0.717 | 0.725 | 0.742 | 0.721 |
| 478590454 | 0.437 | 0.469 | 0.339 | 0.324 | 0.275 |
| 480353286 | 0.554 | 0.604 | 0.537 | 0.613 | 0.583 |
| 502887600 | 0.609 | 0.550 | 0.434 | 0.450 | 0.442 |
| 476595373 | 0.673 | 0.691 | 0.720 | 0.747 | 0.745 |
| 480171386 | 0.600 | 0.743 | 0.634 | 0.582 | 0.522 |
| 485837504 | 0.661 | 0.628 | 0.562 | 0.630 | 0.568 |
| 329549957 | 0.627 | 0.644 | 0.470 | 0.531 | 0.561 |
| 500857324 | 0.548 | 0.570 | 0.356 | 0.555 | 0.584 |
| 324065524 | 0.750 | 0.798 | 0.684 | 0.781 | **0.836** |
| 478498617 | 0.540 | 0.655 | 0.598 | 0.654 | 0.655 |
| 502574847 | 0.726 | 0.717 | 0.831 | 0.828 | 0.828 |
| 313861677 | 0.640 | 0.649 | 0.443 | 0.461 | 0.491 |
| 501951290 | 0.605 | 0.673 | 0.433 | 0.548 | 0.559 |
| 480798873 | 0.700 | 0.663 | 0.702 | 0.725 | 0.730 |
| Cux2 |  |  |  |  |  |
| 487358945 | 0.703 | 0.644 | 0.592 | 0.604 | 0.661 |
| 486239338 | 0.536 | 0.731 | 0.462 | 0.663 | 0.613 |
| 486198953 | 0.692 | 0.645 | 0.619 | 0.764 | 0.765 |
| 490126119 | 0.807 | 0.775 | 0.833 | **0.864** | 0.859 |
| 486108147 | 0.669 | 0.718 | 0.604 | 0.691 | 0.697 |
| Ntsr1 |  |  |  |  |  |
| 488426904 | 0.751 | 0.679 | 0.847 | 0.906 | 0.902 |
| 485339983 | 0.728 | 0.681 | 0.685 | 0.742 | 0.795 |
| 485905411 | 0.608 | 0.567 | 0.795 | 0.783 | 0.777 |
| 471688285 | 0.723 | 0.413 | 0.627 | 0.806 | 0.804 |
| 503814817 | 0.689 | 0.590 | 0.846 | 0.869 | 0.885 |
| 490263438 | 0.840 | 0.723 | 0.842 | 0.887 | **0.910** |
| Scnn1a-Tg3 |  |  |  |  |  |
| 476557750 | 0.669 | 0.663 | 0.639 | 0.732 | 0.735 |
| 486893033 | 0.610 | 0.618 | 0.480 | 0.583 | 0.690 |
| 476263004 | 0.727 | 0.643 | **0.885** | 0.854 | 0.835 |
| 476127518 | 0.614 | 0.689 | 0.592 | 0.743 | 0.746 |
| 476269122 | 0.602 | 0.626 | 0.531 | 0.650 | 0.646 |
| 476271271 | 0.547 | 0.613 | 0.583 | 0.579 | 0.566 |
| Specimen ID | 1 | 2 | 3 | 4 | 5 |
| Nr5a1 |  |  |  |  |  |
| 489731121 | 0.658 | 0.695 | 0.597 | 0.698 | 0.705 |
| 502267531 | 0.587 | 0.621 | 0.529 | 0.666 | 0.654 |
| 502245101 | 0.732 | **0.740** | 0.706 | 0.737 | 0.699 |
| 489750097 | 0.683 | 0.718 | 0.705 | 0.727 | 0.734 |
| 476305247 | 0.691 | 0.693 | 0.653 | 0.716 | 0.732 |
| 502193385 | 0.680 | 0.687 | 0.630 | 0.644 | 0.632 |
| 321893798 | 0.509 | 0.626 | 0.400 | 0.453 | 0.496 |
| 318733871 | 0.570 | 0.704 | 0.472 | 0.711 | 0.732 |
| Htr3a |  |  |  |  |  |
| 482654108 | 0.759 | 0.664 | 0.916 | 0.916 | 0.917 |
| 488686369 | 0.712 | 0.691 | 0.864 | 0.864 | 0.871 |
| 475622793 | 0.694 | 0.656 | 0.917 | 0.905 | **0.919** |
| 482438913 | 0.758 | 0.724 | 0.885 | 0.907 | 0.870 |
| 464188580 | 0.667 | 0.600 | 0.863 | 0.849 | 0.869 |
| 487176969 | 0.851 | 0.871 | 0.885 | 0.887 | 0.890 |
| 519517188 | 0.731 | 0.653 | 0.852 | 0.827 | 0.832 |
| 482516216 | 0.717 | 0.739 | 0.837 | 0.872 | 0.844 |
| 475586130 | 0.678 | 0.663 | 0.790 | 0.783 | 0.769 |
| Sst |  |  |  |  |  |
| 322197295 | 0.420 | 0.397 | 0.406 | 0.483 | 0.491 |
| 485248699 | 0.803 | 0.822 | 0.848 | 0.851 | 0.879 |
| 324495712 | 0.433 | 0.538 | 0.455 | 0.517 | 0.551 |
| 476686112 | 0.694 | 0.747 | 0.709 | 0.684 | 0.760 |
| 328015342 | 0.747 | 0.706 | 0.676 | 0.748 | 0.776 |
| 502382506 | 0.789 | 0.794 | 0.752 | 0.805 | 0.815 |
| 322229594 | 0.671 | 0.753 | 0.640 | 0.682 | 0.662 |
| 504608602 | 0.409 | 0.421 | 0.401 | 0.393 | 0.534 |
| 328031983 | **0.891** | 0.889 | 0.884 | 0.869 | 0.872 |
| 478409545 | 0.823 | 0.837 | 0.845 | 0.858 | 0.852 |
| 475542907 | 0.753 | 0.755 | 0.770 | 0.786 | 0.788 |
| 475057898 | 0.863 | 0.889 | 0.842 | 0.885 | 0.883 |
| Rbp4 |  |  |  |  |  |
| 485577686 | 0.692 | 0.684 | 0.618 | 0.728 | 0.741 |
| 471131311 | 0.775 | 0.737 | 0.688 | 0.754 | 0.736 |
| 490916919 | 0.703 | 0.659 | 0.720 | 0.725 | 0.792 |
| 508911693 | 0.507 | 0.635 | 0.708 | 0.636 | 0.640 |
| 485573974 | 0.720 | 0.717 | 0.725 | 0.785 | 0.776 |
| 490612844 | 0.668 | 0.665 | 0.710 | 0.704 | 0.715 |
| 485835016 | 0.674 | 0.632 | 0.624 | 0.721 | 0.715 |
| 485880739 | 0.618 | 0.614 | 0.663 | 0.665 | 0.606 |
| 490982660 | 0.742 | 0.639 | 0.832 | 0.865 | **0.884** |

2. Z-score all parameters and hierarchically cluster cells using Euclidean distance and Ward’s method.
3. Split cells into two clusters, as defined by the top branch split of the dendrogram from the hierarchical clustering.
4. Calculate the likelihood associated with the data being derived from two multivariate Gaussians (defined by the two clusters) versus a single multivariate Gaussian. This was calculated using the sigClust package in R.
5. If the assessment of the separation into two clusters (from step 4) is significant (sigClust p-value < 0.001), repeat steps 2-4 on each of the two subpopulations.
6. If the assessment of the separation into two clusters (from step 4) is not significant, assign the given population to a terminal cluster.
7. End clustering once all cells are assigned to a terminal cluster that cannot be split further.

Because the estimation of the separation into two clusters (step 4) has some stochasticity (using the current implementation of the sigClust package), clustering was run 100 times for each set of parameters, and only final clusters that were identified in at least 50 runs were included as part of the final run. Given that the identity of cells segregated into two branches is not stochastic (as opposed to the evaluation of the branch separation), all clustering runs yield final trees that only differ in their depth, not in their segregation across branches. As a result, the final set of clusters chosen for a given set of parameters can be coherently assembled into a single tree, with each leaf and intermediate node being assigned a percentage based on the number of runs in which it appeared, as illustrated in the corresponding figure.

### 4.7 Exclusion Criteria

Here, since we are interested in the models of transgenic lines, we include only cre positive cells and and neurons from transgenic lines that have more than 7 modelable neurons. After parameters were fit as described in section “3 Parameter fitting and distributions”, models containing outlier parameters or parameters that are biologically unrealistic as listed below are excluded from the data set.

1. Neurons were eliminated because the training subthreshold noise epochs (used for fitting resistance via linear regression) contained a spike.
2. Neurons with an intercept greater than 30 mV resulting from the linear regression performed during the spike cut width calculation (Figure 8).
3. Neurons with an instantaneous threshold, Θ_∞_, less than −60 mV before post-hoc optimization (Figure 7) are eliminated.
4. Neurons are eliminated because they contain a resistance greater than 1000 MΩ (Figure 9).
5. Neurons that fail to complete the post-hoc optimization without error.

Additional neurons were eliminated from models containing reset rules based on the following criteria:

1. Models had an amplitude of the spike component of threshold, *δ*_*s*_ that was larger than 20 mV (Figure 10).
2. Models had an amplitude of the spike component of threshold, *δ*_*s*_ that was less than zero (Figure 10)
3. Models had 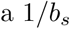 that was greater than 100 mS^−1^ (Figure 10).

Finally models with the following voltage dependent threshold are excluded:

1. Models tha have *a_v_* less than −50 *s*^−1^ (Figure 10).
2. Models that have *a b_v_* is less than 0.1 *s*^−1^ (Figure 10).

This leaves a total of 771 neurons that have a *GLIF*_1_, and *GLIF*_3_ model, 289 neurons that have a *GLIF*_2_ and a *GLIF*_4_ model, and 288 neurons that have a *GLIF*_5_ model.

### 4.8 Using linear regression to solve for variables using a noisy stimulus

One way to fit the resistance, *R*, capacitance, *C*, and resting potential, *El* is to use linear regression on the voltage, *V*, response to a noisy stimulus, *I*. The standard leaky integrate and fire model for the dynamics of a neuron is:

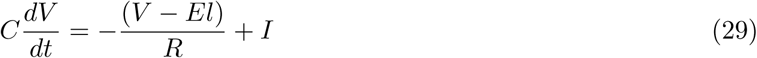

This equation can be rearranged so that it fits into the form *y* = *Xβ* + *∈* where *y* is the dependent variable, *X* is a the independent variable and are the parameters being fit. For implementation this means that the parameters being fit should be associated with the matrix on the (rhs) of the equation, *lhs* and the known parameters should be on the left hand side (lhs) of the equation After the equation has been arranged in the proper form, a standard least squares algorithm such as *np:linalg:lstsq* in python can be used.

For example let us say we want tot El and R in Equation 29. Equation 29 can be rearranges as follows:

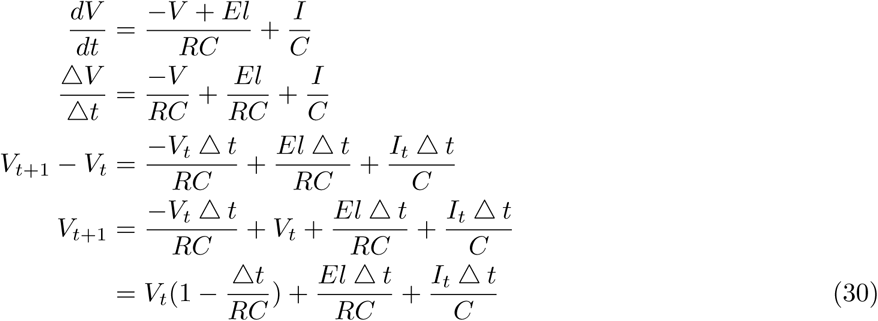

Here, Δ*t* is the amount of time between *V* and *V*_*t*+1_ (time step) where *t* is the timestep index which runs from *t* = 1 to *t* = *T* where *T* is the total number of time steps in the data.

For implementation, each term must be associated with a vector. The vector may be output data or input stimulus (for example, here *V*, *V*_*t*+1_, or *I*) or a vector of ones when a term is not associated with a time series. In vector form, Equation 30 is:

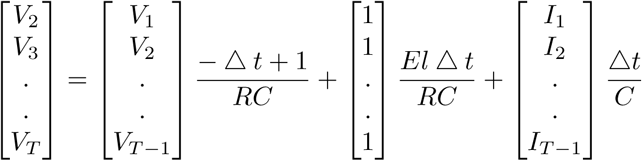

Fitting R, C and El results in the following equation:

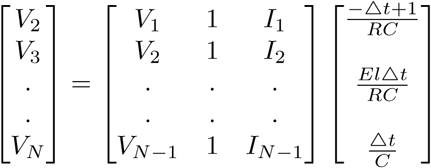

The output will be an vector where

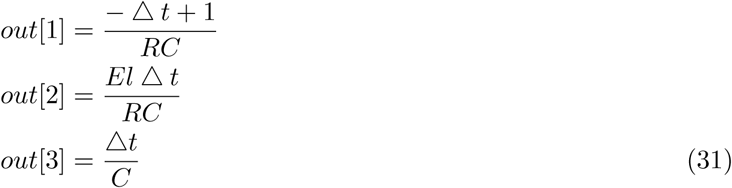

Solving this system of equations yields:

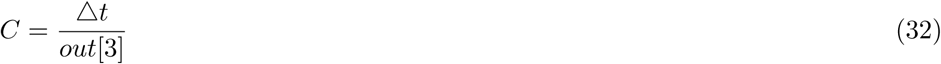

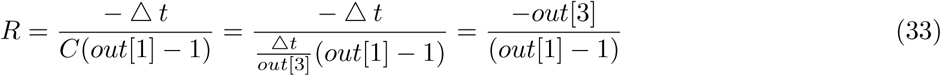

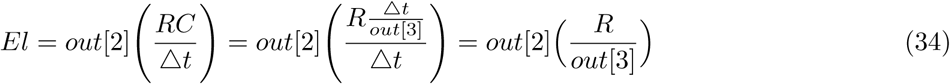

Now let’s say that we do not want to fit the capacitance during the regression. In this case the term containing the *I* vector will be shifted to the left hand side of the equation:

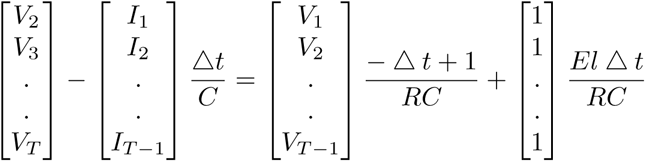

In matrix form:

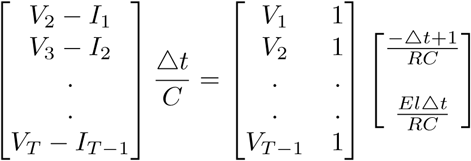

Solving this system of equations yields:

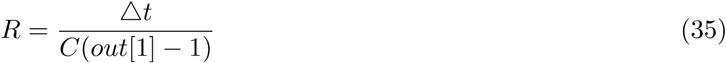

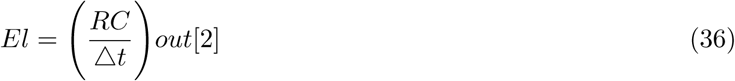

Above, we solved this problem by separating Δ*V* into the voltages before and after the time step Δt, i.e. *V*_n+1_ - *V*. This problem can also be solved by calculating 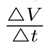 directly from the data.

### 4.9 Analytical solution for the dynamics of the voltage component of the threshold

The voltage component of the threshold evolves according to the following dynamics:

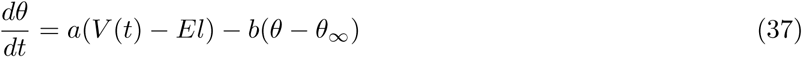

Where θ is the voltage component of the threshold, *V* is the voltage of the neuron which changes in time *t*, *El* is the resting potential, Θ_∞_, and *a* and *b* are constants which will be fit to the data.

Below we willnd the analytical solution to equation (37).

The solution to a di erential equation with the form:

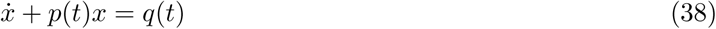

is

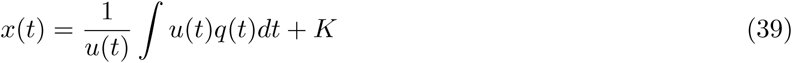

where *K* is the standard integration constant and

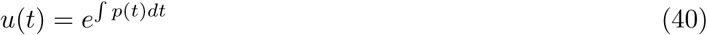

Equation (37) can be rearranged into this form:

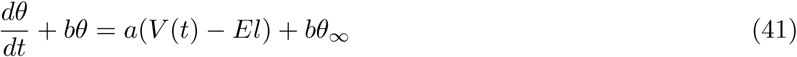

Where

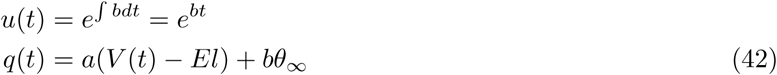

Therefore:

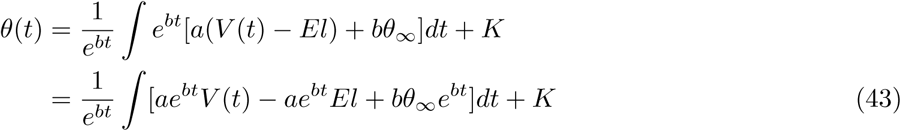

The analytical solution for *V*(*t*)

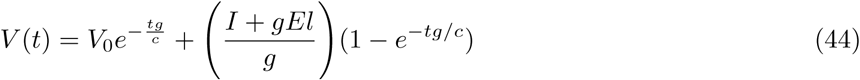

Plugging in *V*(*t*) for θ(*t*) and defining 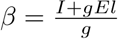 gives:

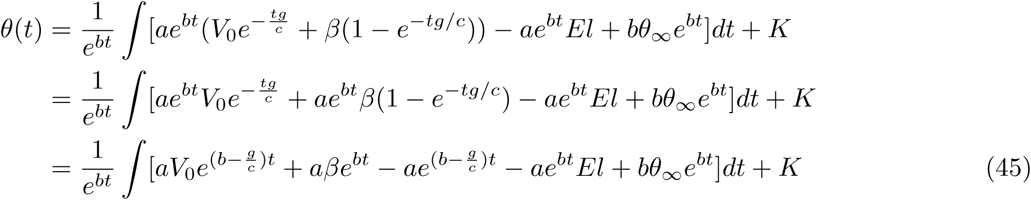

Setting 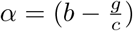

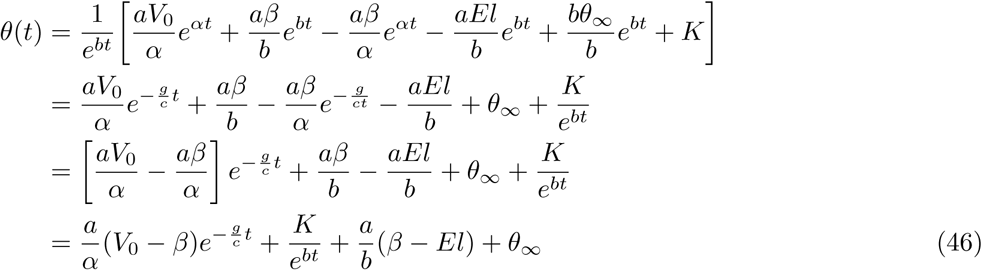

at *t* = 0, θ(*t*) = θ_0_. Therefore:

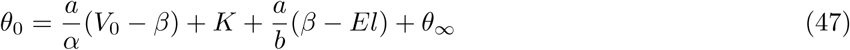

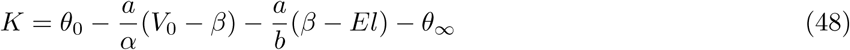

Plugging in *K*:

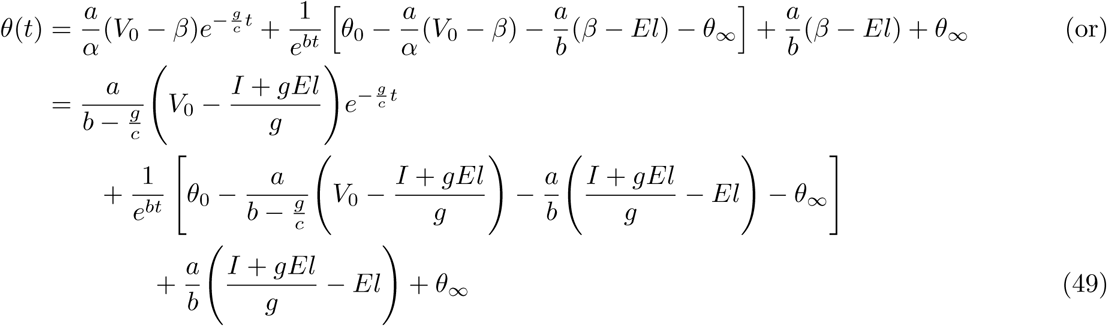

**Table 6:** Summary characterization of GLIF parameters. All Cre lines with 7 or more LIF neurons are included. The single number in each cell is the median. The low and high quartiles are edspeci in brackets below the median. Some neurons have the required stimuli for LIF models but do not have the stimuli required for higher level models. When there are not more than 7 neurons that have the parameter, these values are denoted with a NaN.

| | R (MΩ) | $\tau$ (ms) | C (pF) | El (mV) | th$_{\infty}$ (mV) | $\Delta$ th$_{\infty}$ (mV) | R$_{ASC}$ (MΩ) | Q$_1$ (pC) |
| --- | --- | --- | --- | --- | --- | --- | --- | --- |
| inhibitory | 193 | 11.4 | 59.7 | -72.4 | -45.4 | 26.6 | 201 | -1.28 |
|  | [135, 278] | [7.98, 17.9] | [50.6, 71.9] | [-75.3, -69.2] | [-47.6, -42.8] | [23.8, 30.2] | [140, 309] | [-2.32, -0.742] |
| excitatory | 180 | 18.7 | 106 | -75.3 | -47 | 27.6 | 191 | -2.84 |
|  | [136, 244] | [14.7, 27] | [88.6, 129] | [-78.1, -71.8] | [-48.9, -45.1] | [24.9, 31] | [138, 280] | [-3.96, -2.04] |
| Scnn1a-Tg2 | 179 | 27.7 | 129 | -74.4 | -46 | 26.8 | 215 | -3.36 |
|  | [126, 232] | [16.9, 31.7] | [112, 166] | [-76.4, -71.9] | [-49.5, -44.7] | [25.5, 29.8] | [126, 321] | [-4.18, -2.13] |
| Nr5a1 | 188 | 17.7 | 94.9 | -77 | -47.2 | 30.8 | 184 | -2.65 |
|  | [146, 265] | [13.8, 24.5] | [83.3, 107] | [-80.2, -73.3] | [-49.3, -45.4] | [27.4, 32.1] | [129, 265] | [-3.54, -2.21] |
| Scnn1a-Tg3 | 189 | 20.9 | 104 | -75.7 | -46.4 | 28.6 | 212 | -2.54 |
|  | [154, 246] | [14.4, 28.2] | [86.6, 131] | [-78, -72] | [-48.3, -44.6] | [25, 32.8] | [146, 294] | [-3.58, -1.68] |
| Rorb | 192 | 19.1 | 98.3 | -75.2 | -46.8 | 28 | 197 | -2.78 |
|  | [143, 262] | [15.4, 29.1] | [86.9, 123] | [-78.3, -71.3] | [-49.1, -44.9] | [25.2, 30.8] | [150, 299] | [-3.76, -1.99] |
| Cux2 | 166 | 16.8 | 105 | -75.3 | -47.2 | 27.5 | 175 | -2.69 |
|  | [136, 221] | [14, 21.5] | [88.5, 118] | [-78.1, -72.7] | [-48.4, -45.5] | [26, 30.9] | [137, 223] | [-3.94, -1.91] |
| Ntsr1 | 186 | 16.7 | 91.2 | -75.4 | -47.3 | 28.1 | 192 | -2.9 |
|  | [153, 227] | [15, 19.3] | [80.5, 104] | [-77.7, -72.3] | [-48.3, -45.4] | [25.6, 30.5] | [146, 232] | [-4.75, -2.13] |
| Ctgf | 245 | 25.5 | 119 | -73.7 | -46.3 | 24.5 | 275 | -2.36 |
|  | [160, 345] | [20.5, 37] | [97, 135] | [-75.4, -67.6] | [-48.5, -44.7] | [19.2, 28.2] | [183, 405] | [-3.13, -1.83] |
| Rbp4 | 137 | 18.3 | 135 | -73.8 | -47.8 | 24.9 | 162 | -3.18 |
|  | [104, 195] | [14, 26] | [111, 150] | [-77.8, -68.8] | [-49.4, -45.9] | [22.2, 28.9] | [113, 253] | [-4.54, -2.22] |
| Sst | 234 | 16.7 | 70 | -71.3 | -45.8 | 25.3 | 242 | -1.36 |
|  | [171, 319] | [11.2, 22.1] | [57.7, 84.2] | [-73.9, -68.4] | [-47.9, -43.4] | [22.3, 28.8] | [168, 340] | [-1.78, -0.858] |
| Pvalb | 137 | 7.55 | 54.4 | -74.6 | -46.2 | 27.6 | 143 | -0.734 |
|  | [110, 179] | [6.52, 9.42] | [49.5, 65.3] | [-76.4, -71.5] | [-48, -43.6] | [24.9, 30.5] | [109, 180] | [-1.24, -0.507] |
| Htr3a | 195 | 11.1 | 55.4 | -72.1 | -44 | 26.9 | 207 | -2.07 |
|  | [136, 335] | [7.92, 16.4] | [45.2, 66.2] | [-75, -67.7] | [-45.9, -41] | [24, 30.8] | [148, 345] | [-3.84, -1.02] |
| Ndnf | 189 | 11.9 | 62.3 | -72.7 | -42.4 | 29.7 | 203 | -2.32 |
|  | [140, 215] | [11.5, 13] | [59.4, 74.5] | [-75.3, -68.3] | [-42.8, -40.6] | [26.4, 35.1] | [154, 234] | [-2.56, -1.93] |
| Chat | 223 | 14.9 | 55.3 | -74.1 | -47.3 | 26.6 | 226 | -1.25 |
|  | [181, 325] | [11, 20] | [50.9, 64.3] | [-76.8, -73] | [-47.7, -46.7] | [25.6, 28.8] | [189, 338] | [-1.85, -0.895] |
| Vip | 324 | 20.3 | 57.9 | -72.2 | -46.3 | 25.1 | 342 | -1.04 |
|  | [260, 422] | [14.2, 23.2] | [51.5, 66.2] | [-74.6, -69.8] | [-48, -44.7] | [22.7, 27.4] | [277, 492] | [-2.28, -0.807] |

**Table 7:**
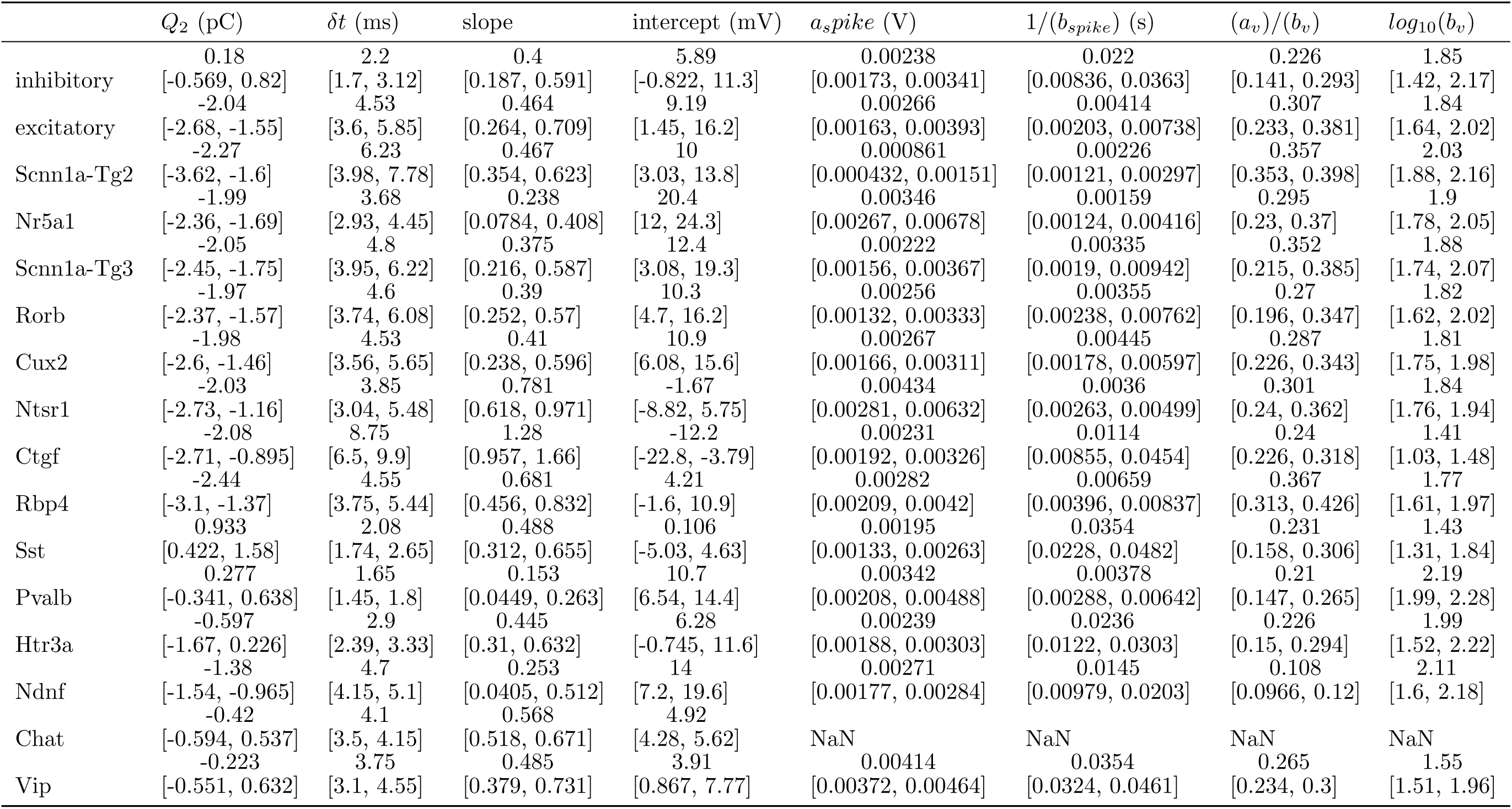
SECOND HALF OF TABLE 6.

**Table 8:**
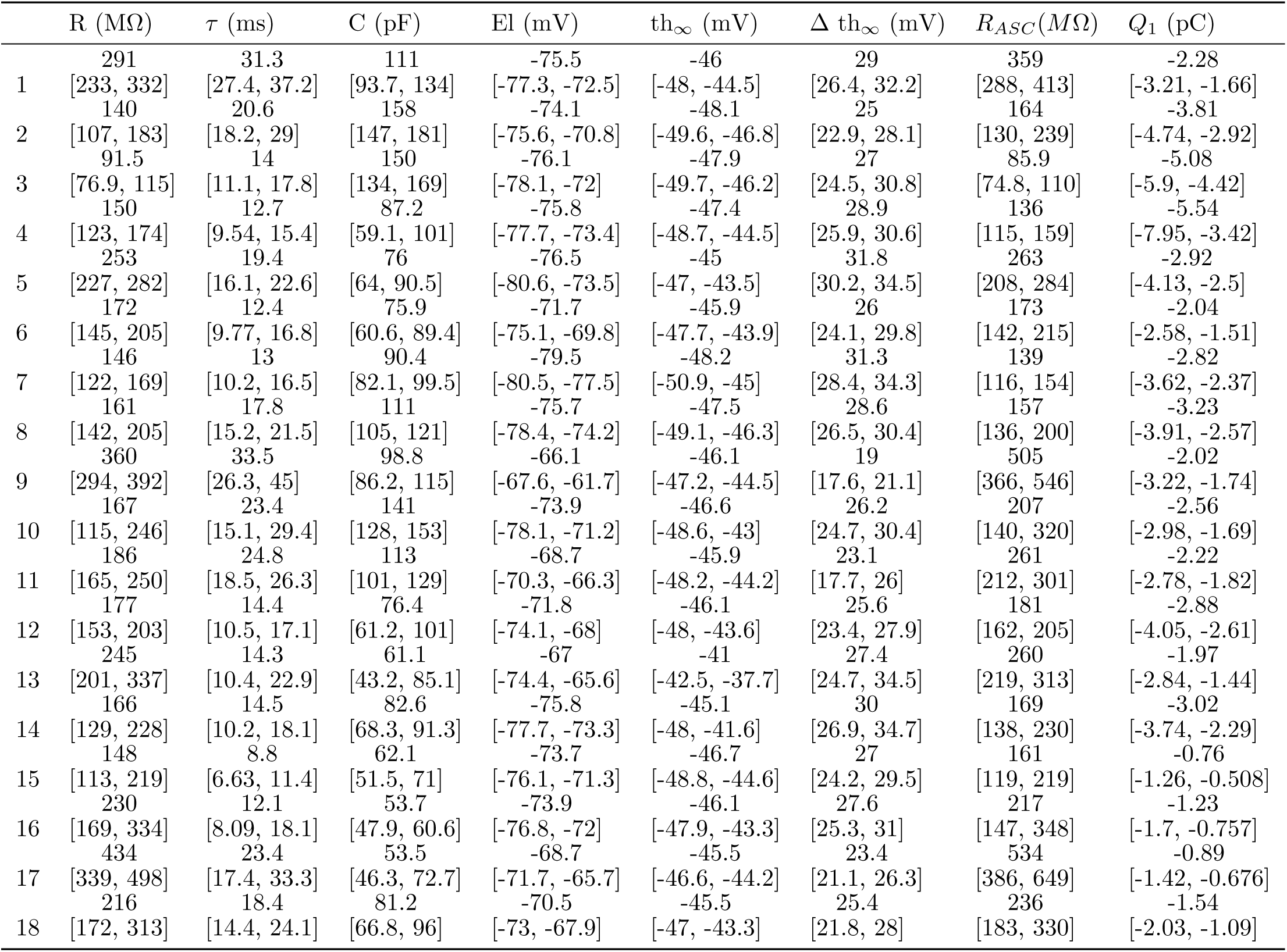
Summary characterization of GLIF parameters within clusters. The single number in each cell is the median. The low and high quartiles are demarked in brackets below the median. Some neurons have the required stimuli for LIF models but do not have the stimuli required for higher level models. When there are less than 7 neurons that have the parameter, these values are denoted with a NaN.

**Table 9:**
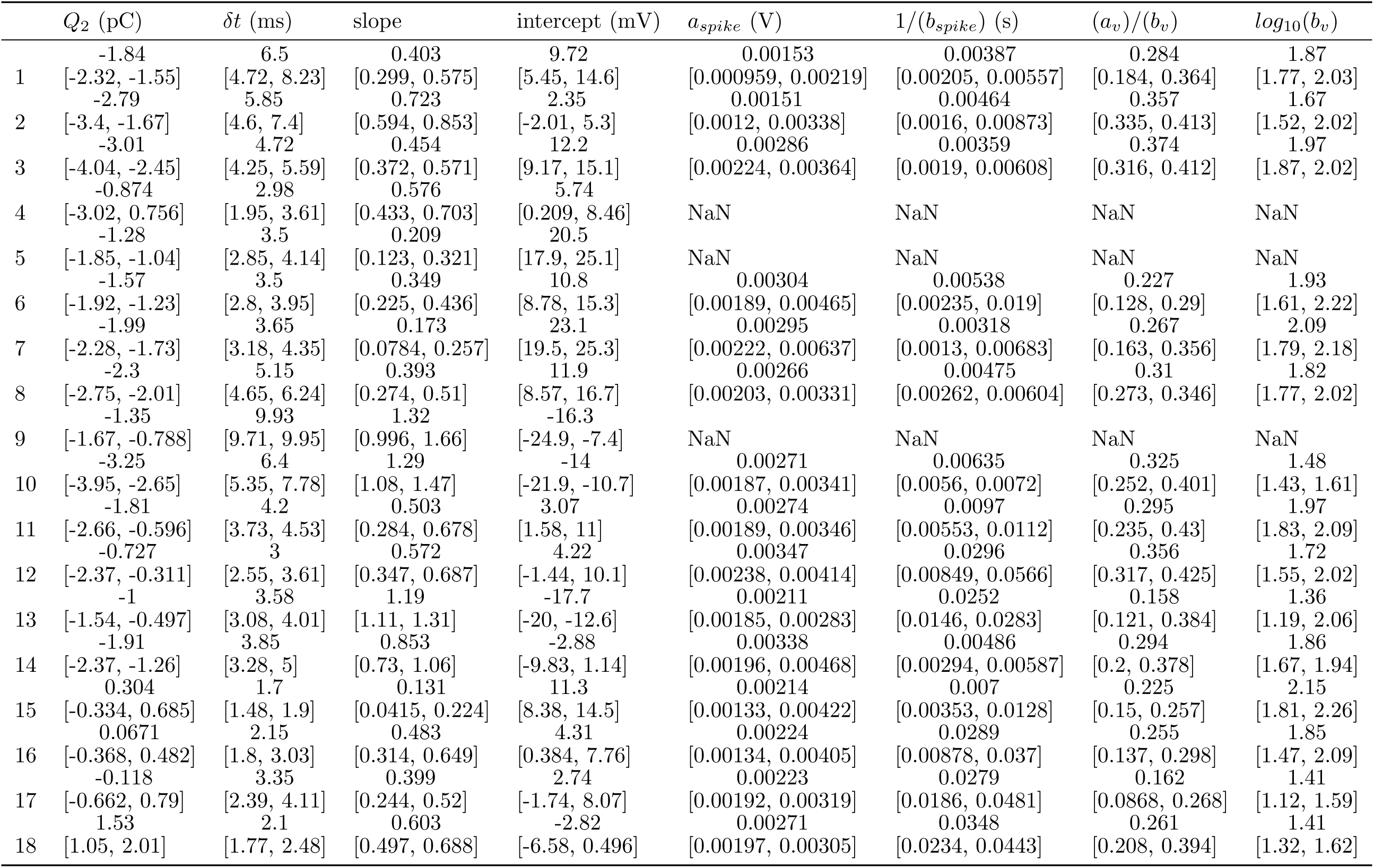
SECOND HALF OF TABLE 8.

**Table 10:** Data features of clusters found by clustering GLIF parameters: * short square, † long square.

| | $\tau_m (ms)$ | $R_i (M\Omega)$ | $V_{rest} (mV)$ | $I_{threshold}^\dagger (pA)$ | $V_{threshold}^\dagger (mV)$ | $V_{peak}^\dagger (mV)$ | $V_{fasttrough}^\dagger (mV)$ | $V_{trough}^\dagger$ |
| --- | --- | --- | --- | --- | --- | --- | --- | --- |
| 1 | 24.3<br>[20.3, 28.8] | 188<br>[148, 218] | -75<br>[-77.2, -72.2] | 70<br>[50, 90] | -36.8<br>[-39.1, -35.2] | 38.5<br>[33.7, 44.4] | -48<br>[-50.4, -45.9] | -55.7<br>[-58.4, -54.1] |
| 2 | 20<br>[15.5, 23.6] | 125<br>[103, 144] | -73.9<br>[-75.4, -70.9] | 110<br>[90, 150] | -38.2<br>[-41.4, -36] | 36.6<br>[32.8, 41.2] | -48.4<br>[-49.9, -44.4] | -55.9<br>[-58.3, -54.1] |
| 3 | 18.2<br>[15.4, 20.2] | 103<br>[83.6, 127] | -76.1<br>[-78, -71.9] | 210<br>[160, 270] | -36.5<br>[-38.4, -34.3] | 36.7<br>[33.2, 40.4] | -47<br>[-48.2, -45.4] | -53.3<br>[-55.3, -50.7] |
| 4 | 12.3<br>[8.88, 15.3] | 137<br>[110, 161] | -75.8<br>[-77.7, -73.1] | 245<br>[210, 325] | -34.6<br>[-39.5, -32] | 26.6<br>[14.6, 32.7] | -50.2<br>[-59.2, -46.9] | -52.5<br>[-59.3, -48.1] |
| 5 | 19.7<br>[14.4, 22.3] | 189<br>[170, 212] | -76.1<br>[-80.5, -73.2] | 110<br>[87.5, 122] | -35.2<br>[-39.9, -31.6] | 33.8<br>[29.6, 41.1] | -48.6<br>[-52.2, -45.9] | -52.9<br>[-55.7, -49.8] |
| 6 | 16.9<br>[10, 21.1] | 175<br>[146, 215] | -71.7<br>[-75.3, -70.2] | 90<br>[70, 140] | -38.3<br>[-41.2, -36.2] | 37.3<br>[26.5, 43] | -50.7<br>[-52.6, -48.7] | -54.4<br>[-56.5, -52.8] |
| 7 | 17.8<br>[13.9, 20.5] | 149<br>[125, 171] | -79<br>[-80.7, -77.2] | 150<br>[110, 190] | -37<br>[-39.8, -35.8] | 38.9<br>[32.2, 40.9] | -50.6<br>[-53.3, -48.5] | -54.8<br>[-56.6, -52.5] |
| 8 | 20.7<br>[18.4, 23.1] | 164<br>[151, 190] | -75.6<br>[-78.3, -74] | 110<br>[90, 130] | -36.3<br>[-38.6, -33.4] | 37.6<br>[35, 39.7] | -47.6<br>[-49.1, -46] | -53.9<br>[-55.6, -51.9] |
| 9 | 23.2<br>[18.8, 26.7] | 213<br>[207, 256] | -65.8<br>[-67.1, -61.4] | 45<br>[32.5, 50] | -41.2<br>[-43.1, -40.3] | 38.1<br>[35, 39.5] | -50.6<br>[-51, -47.9] | -55.9<br>[-57.7, -55] |
| 10 | 18.7<br>[14.2, 24.8] | 150<br>[91.1, 163] | -74.2<br>[-77.6, -71] | 110<br>[90, 190] | -39.8<br>[-42.8, -35.6] | 38.7<br>[35.8, 41.9] | -46<br>[-49, -44.5] | -54.5<br>[-56, -53] |
| 11 | 18.7<br>[16.4, 25.1] | 171<br>[132, 213] | -68.4<br>[-70.1, -66.5] | 70<br>[55, 90] | -39.8<br>[-41.7, -38.2] | 36.8<br>[33.9, 42.7] | -48.8<br>[-51.9, -46.7] | -56.2<br>[-57.3, -54.1] |
| 12 | 14.2<br>[10.2, 18.3] | 148<br>[130, 178] | -71.3<br>[-74.1, -68.3] | 130<br>[110, 150] | -38<br>[-41.4, -34.8] | 31.2<br>[23, 37.1] | -51.3<br>[-53.8, -49.3] | -54.3<br>[-56.1, -52.2] |
| 13 | 20.1<br>[10.1, 23.6] | 219<br>[187, 254] | -66.9<br>[-73.3, -65.7] | 85<br>[70, 120] | -34.6<br>[-38.4, -32.3] | 26.9<br>[17.6, 30.7] | -49.1<br>[-51.5, -46.9] | -51.6<br>[-55.2, -48.8] |
| 14 | 14.2<br>[11.8, 19.3] | 152<br>[119, 196] | -75.3<br>[-77.4, -73] | 150<br>[100, 220] | -35.8<br>[-38.1, -33.2] | 33.3<br>[26.6, 37.9] | -47.9<br>[-50.6, -46] | -51.7<br>[-54.2, -48.8] |
| 15 | 7.01<br>[6.1, 10.3] | 114<br>[82.7, 149] | -73.3<br>[-75.9, -70.8] | 190<br>[130, 290] | -39<br>[-42.7, -35.8] | 16.5<br>[12.5, 25] | -60.4<br>[-64.2, -56.6] | -60.6<br>[-64.4, -56.9] |
| 16 | 12<br>[7.11, 15.1] | 180<br>[125, 243] | -74<br>[-76.5, -71.9] | 130<br>[72.5, 228] | -39.1<br>[-43.8, -36.1] | 20.3<br>[13.5, 29.9] | -56.8<br>[-62.1, -52.5] | -57<br>[-62.2, -53.7] |
| 17 | 19.7<br>[14.5, 26.1] | 279<br>[240, 339] | -67.9<br>[-72, -65.3] | 30<br>[30, 50] | -43<br>[-45.7, -39.1] | 28.8<br>[22.3, 35.3] | -55.1<br>[-60.9, -50.8] | -57.5<br>[-61.5, -54.1] |
| 18 | 19.7<br>[14.9, 26.7] | 205<br>[170, 249] | -70.6<br>[-72.6, -67.1] | 90<br>[52.5, 110] | -42.2<br>[-45.2, -40.1] | 28.7<br>[22.5, 34.2] | -58.4<br>[-60.5, -55.8] | -58.5<br>[-60.6, -56] |

**Table 11:**
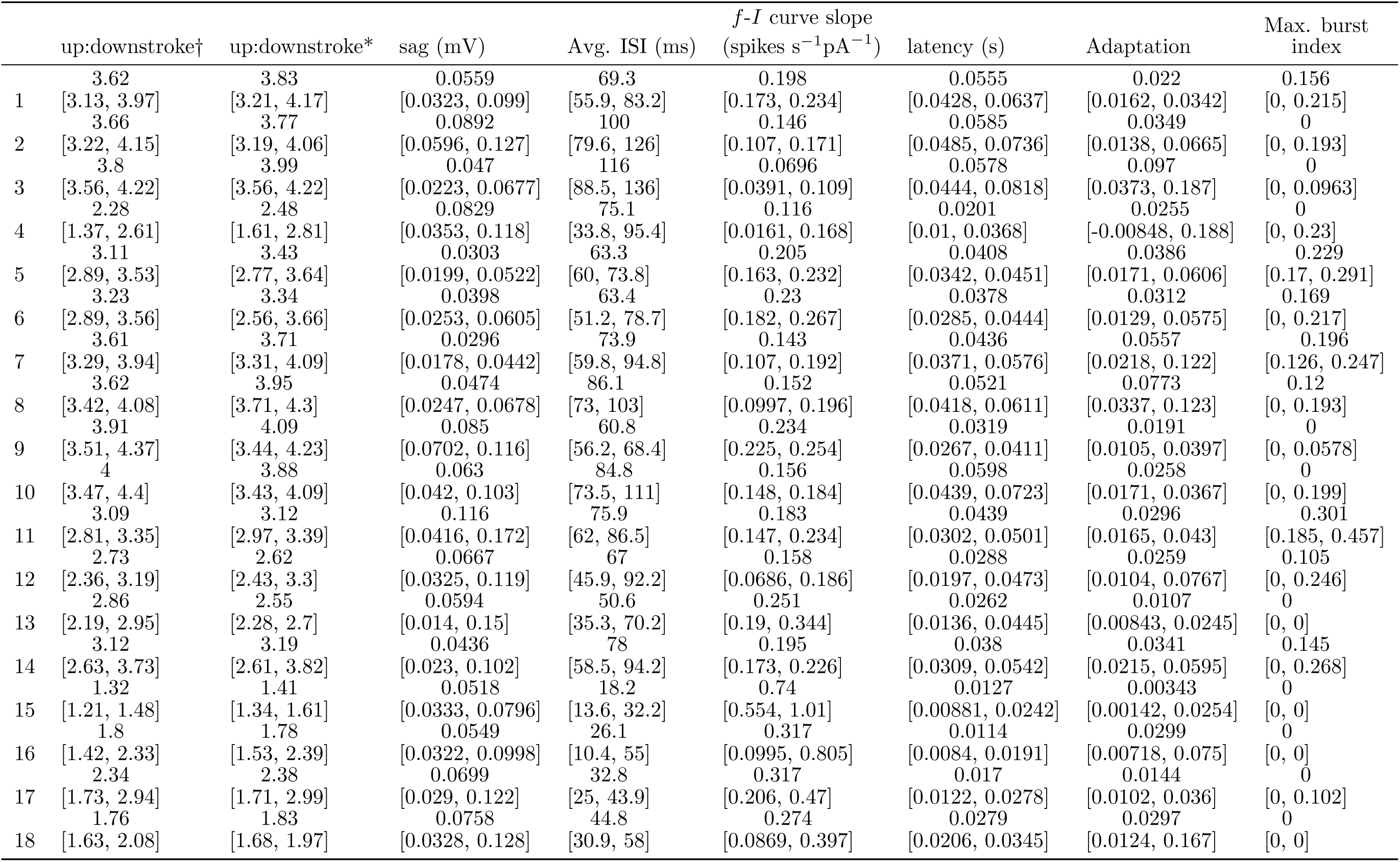
Second half of Table 10.

**Figure 9:**
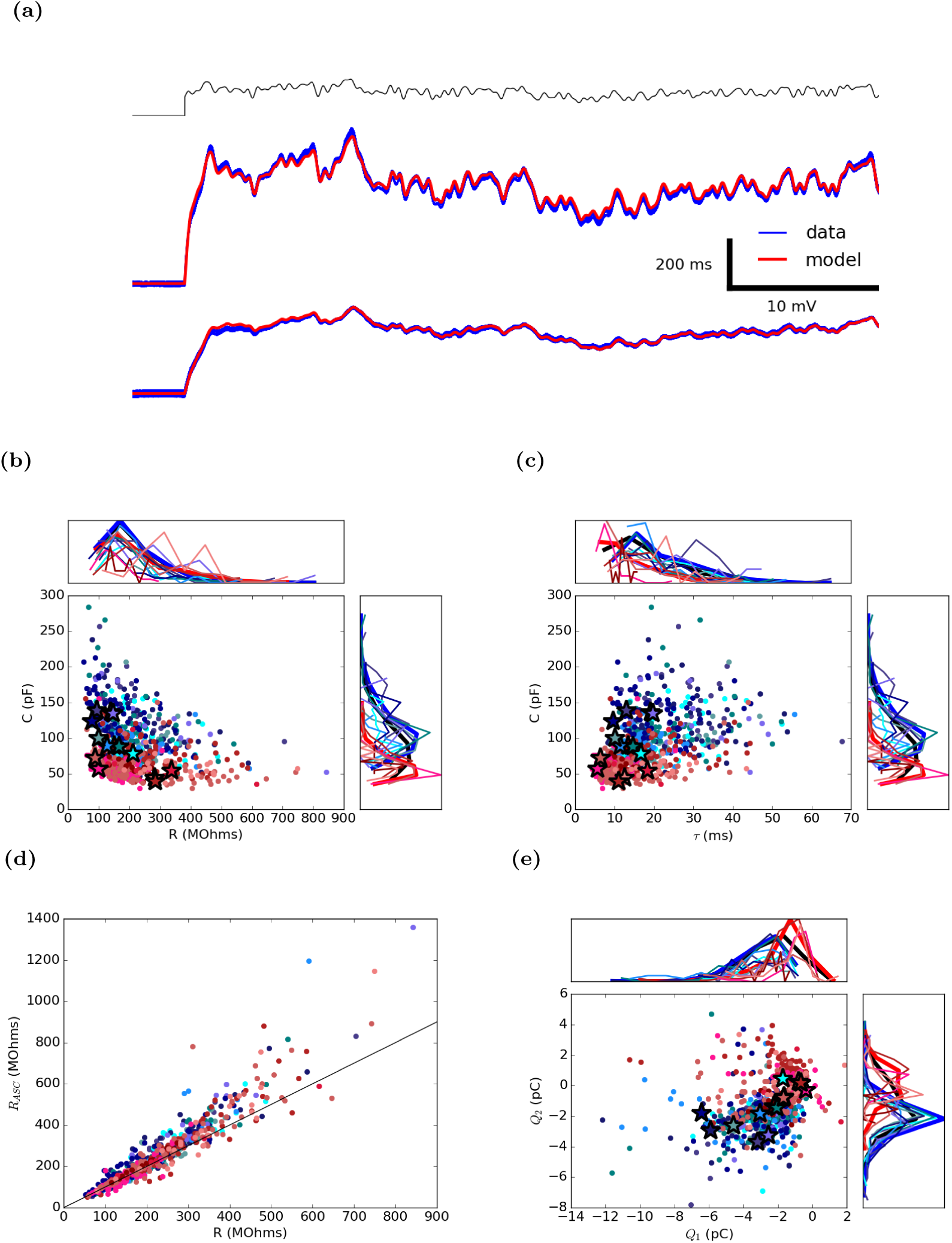
Resistance, Capacitance and amplitudes of after-spike currents are fit via linear regression. (a) Two examples of subthreshold voltage epochs fit via linear regression. Top traces are from the example cells shown in all figure examples (Top: Htr3a 477975366, Bottom: Rorb 314822529). Injected current trace shown in black, voltage of biological neuron recorded from repeated current injections are in blue, voltage of model neuron red. Values of (b) resistance and capacitance and (c) the membrane time constant τ =R*C and capacitance of all neurons fit to subthreshold data without the simultaneous fitting of after-spike currents. (d) Comparison of resistance values obtained by fitting subthreshold data without after-spike currents and supra-threshold data with the simultaneous tting of after-spike currents. (e) Total charge deposited by after-spike currents. In scatter plots and corresponding distributions, shades of blue represent excitatory transgenic lines and red denote inhibitory transgenic lines. The thick black histograms represent data from all transgenic lines together, the thick red histograms represent data from inhibitory transgenic lines and the thick blue histogram represents data from excitatory transgenic lines. Stars denote canonical neurons. The full list of colors corresponding to specific transgenic lines can be found in Table 2 Transgenic Lines. Any neuron containing a spike in the subthreshold noise epoch of the training set was globally eliminated from the data set. (c) and (e) are shown in main article (Figure 3)

**Figure 10:**
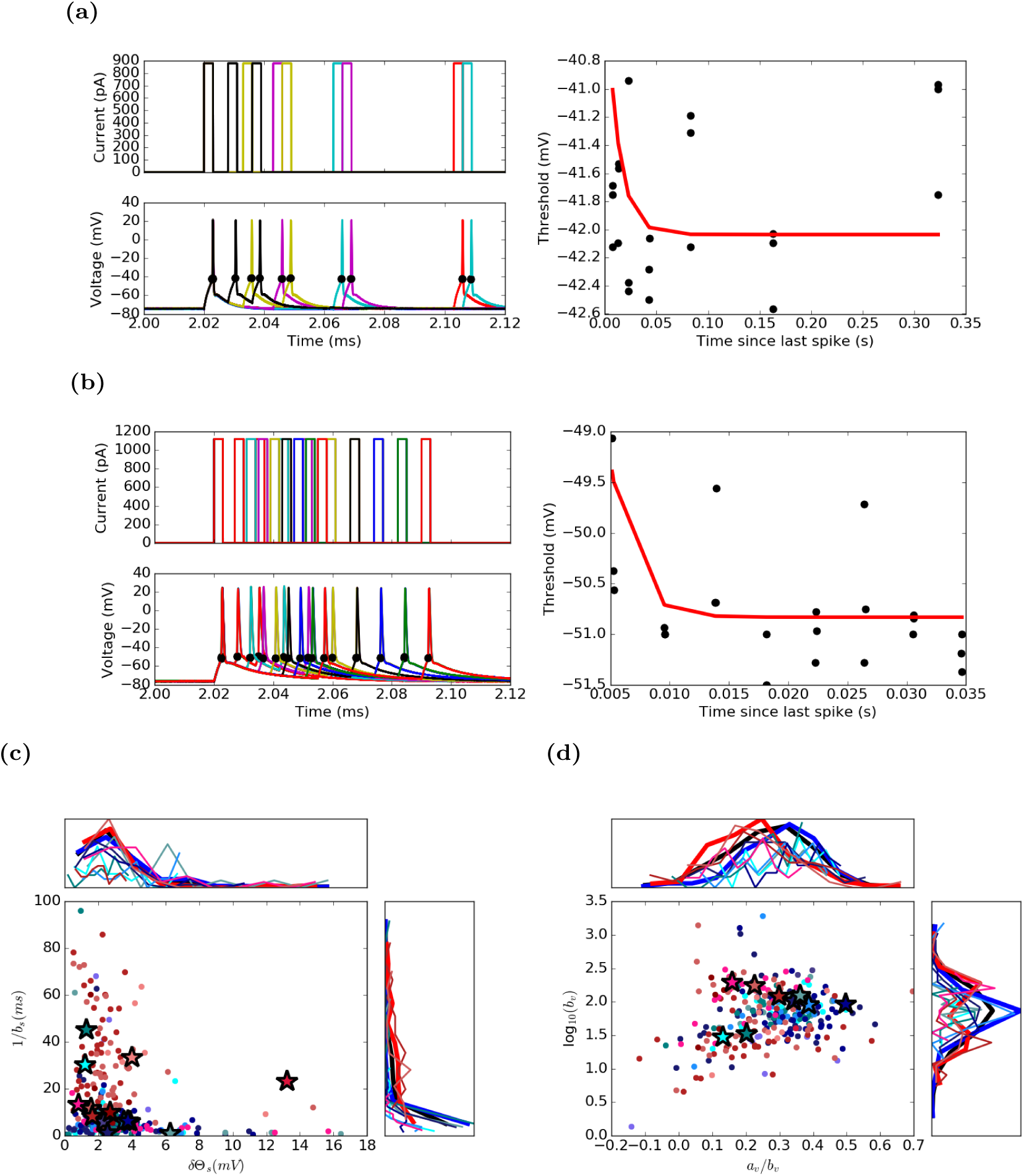
The spiking component and voltage component of the threshold are incorporated into higher level models. The spiking component of the threshold is defined by an exponential function fit to the time between spikes and the voltage at spike initiation of the triple short square data set. Exponential fitting of (a) Htr3a 477975366 and (b) Rorb 314822529. Spiking in response to current pulses (left, with colors representing different sweeps) and exponential fit (right). Black dots denote spike initiation. (c) shows the fit amplitude, *a*, incorporated by the reset rule *δ_s_* and the decay constant, *b* of all neurons with triple square pulse data. All neurons with an 0 < a < 20*mV* or a 1/b > 0.1s are excluded. (d) In the *GLIF*_5_ model the threshold is inuenced by the voltage of the neuron. Parameters from equation 20 are plotted. *GLIF*_5_ models where *a_v_* is < −50 or *b_v_* is < 0.1 are excluded. (c) and (d) are shown in main article (Figure 3)

**Figure 11:**
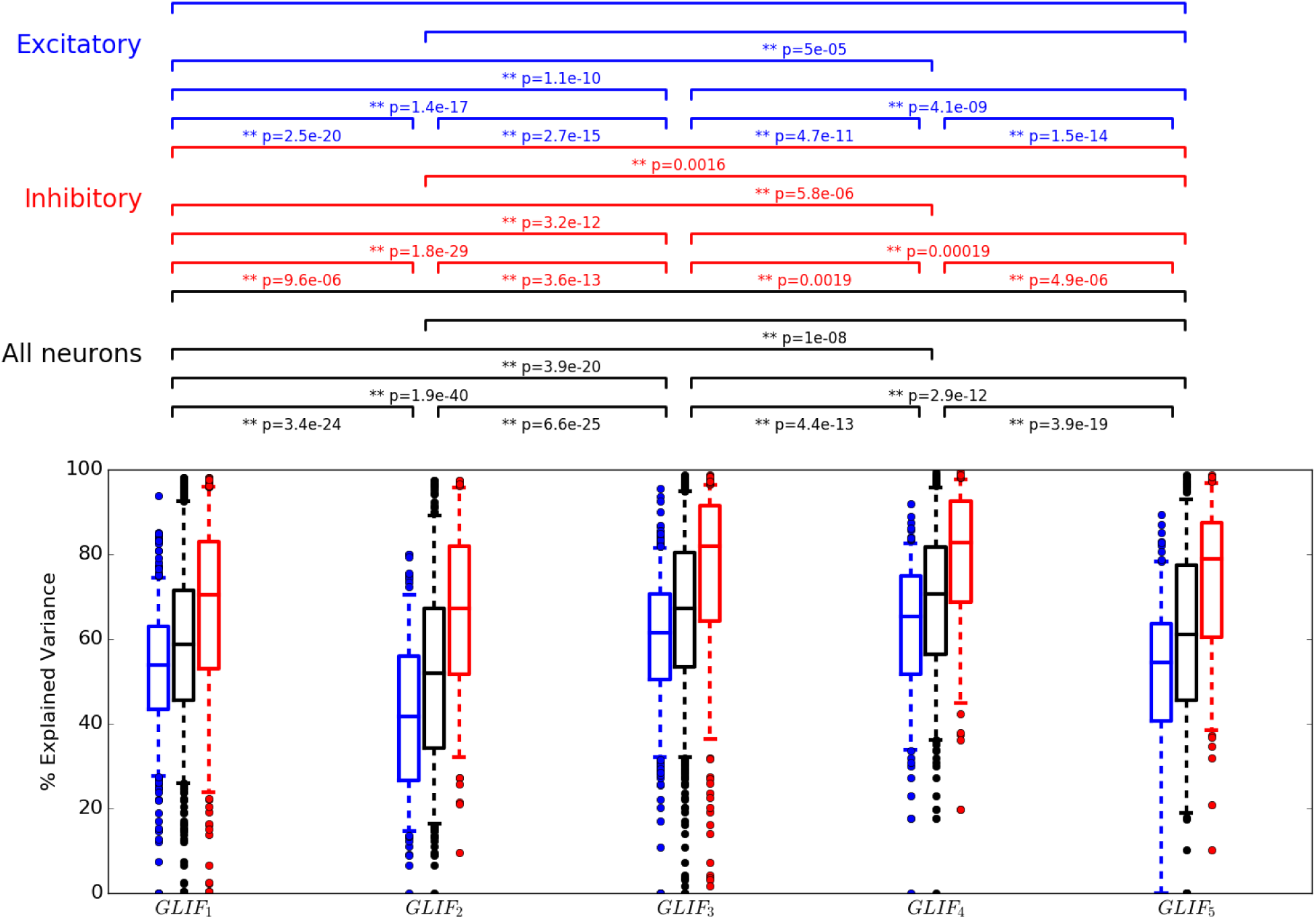
Post-hoc Optimization of *θ*_∞_ improves explained variance. Summary for *GLIF* models for all 771 neurons (black), all 315 inhibitory neurons (red) and all 456 excitatory neurons (blue) for models with measured as opposed to post-hoc optimized *θ*_∞_. Although on average *θ*_∞_ values change very little (Figure 7) before and after optimization, average explained variance values rise (Can an be seen by comparing this figure with Figure 5 of the main text). Box plots display median, and quartiles. Whiskers reach to 5% and 95% quartiles. Individual data points lie outside whiskers. Brackets above plot denote significant difference between distributions assessed via a Wilcoxon sign ranked test: a single asterisk (*) represents a p-value < 0.05, a double asterisk (**) represents a p-value < 0.01.

**Figure 12:**
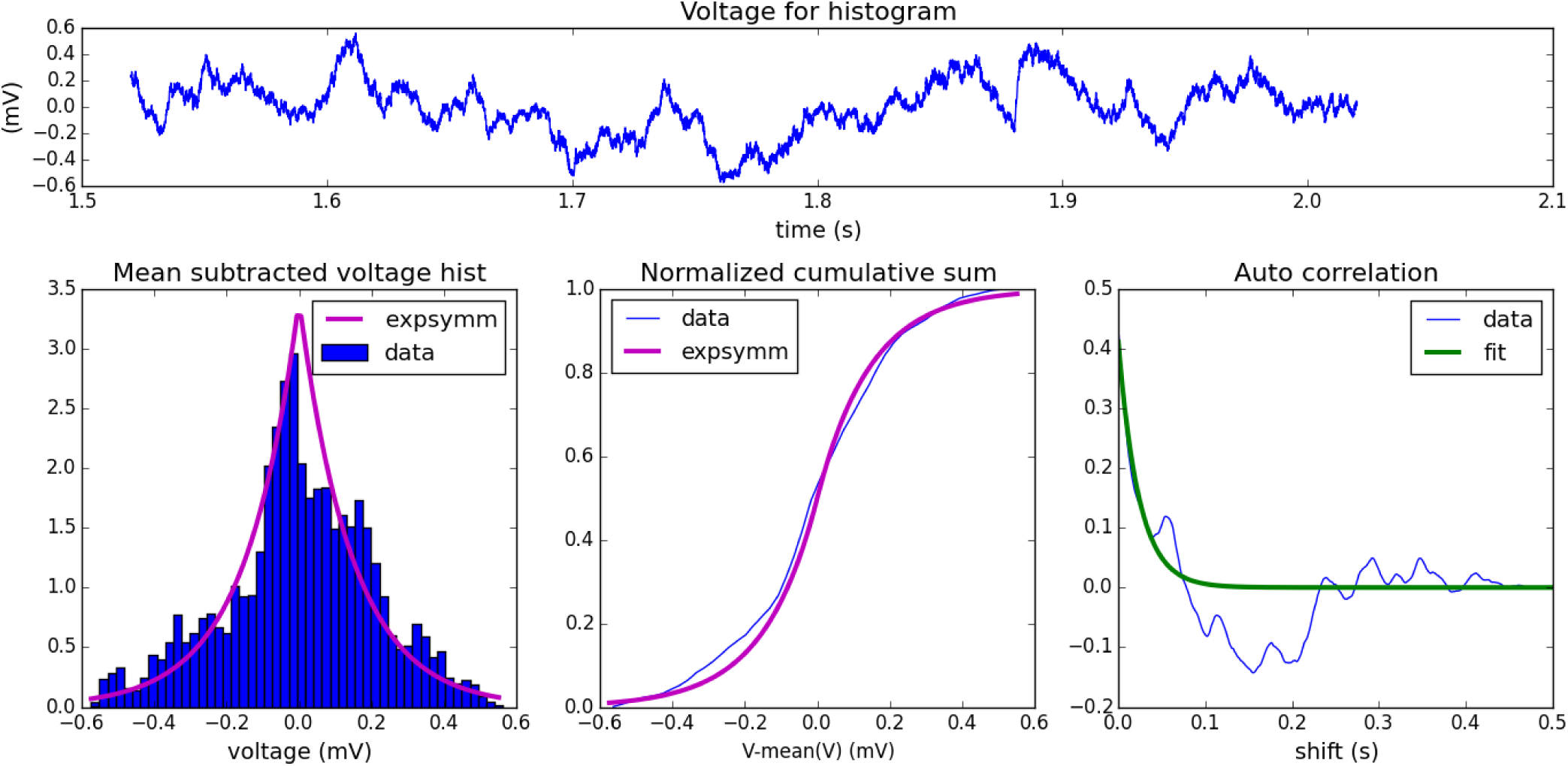
Parameters of the MLIN objective function were extracted from the data. A distribution of voltages is created from the tailing end of the largest amplitude subthreshold square pulse available (top panel). This distribution is then fit by a symmetrically decaying exponential function denoted as expsymm in the plot legends. The width of the non-spiking bins is chosen via a fit of the autocorrelation.

**Figure 13:**
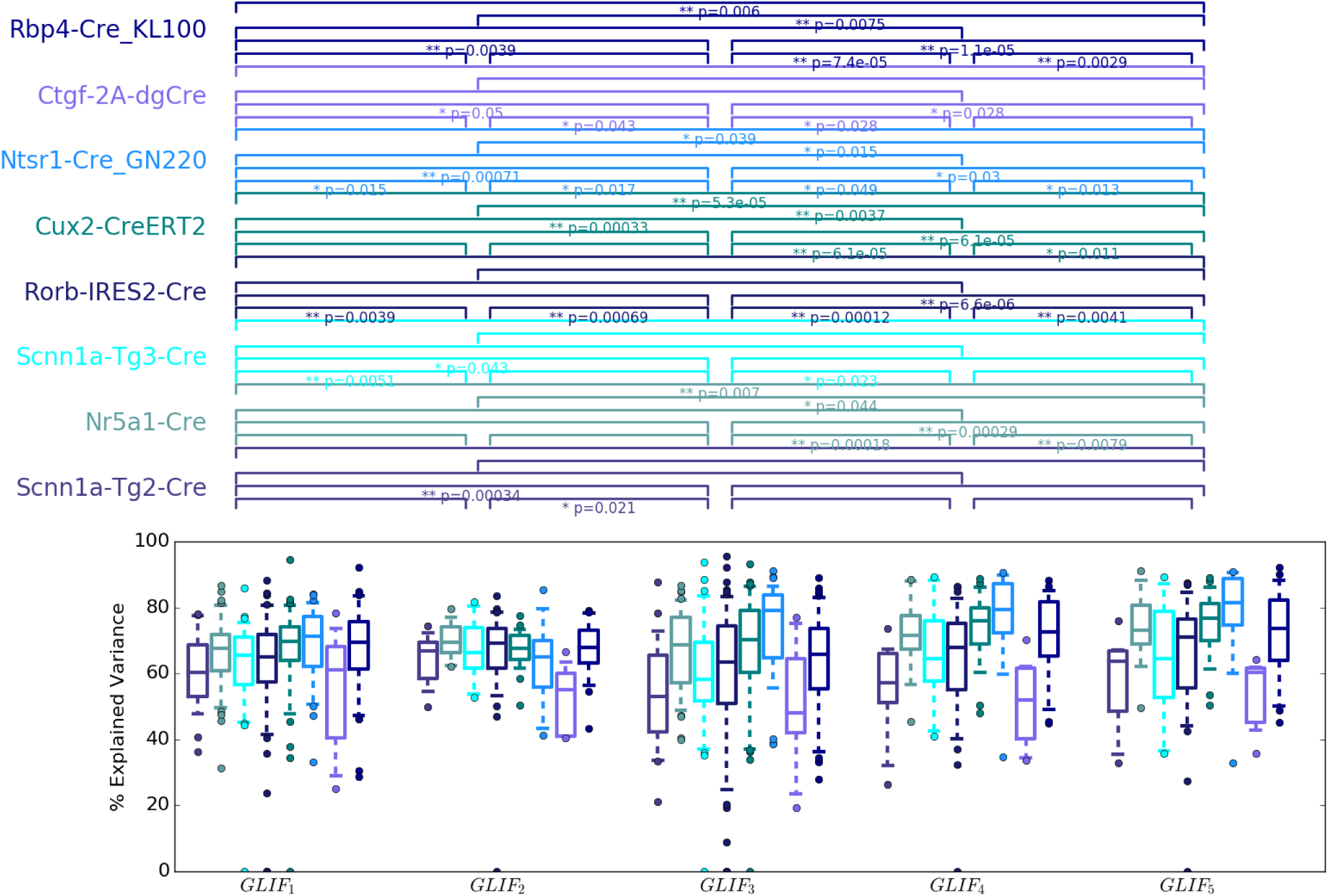
Explained variance summary for excitatory GLIF models of different levels using threshold infinity obtained via MLIN optimization. Box plots display median, and quartiles. Whiskers reach to 5% and 95% quartiles. Individual data points lie outside whiskers. Brackets above plot denote signi cant difference between distributions assessed via a Wilcoxon sign ranked test: a single asterisk (*) represents a p-value < 0.05, a double asterisk (**) represents a p-value < 0.01. Distribution values are available in Table 4

**Figure 14:**
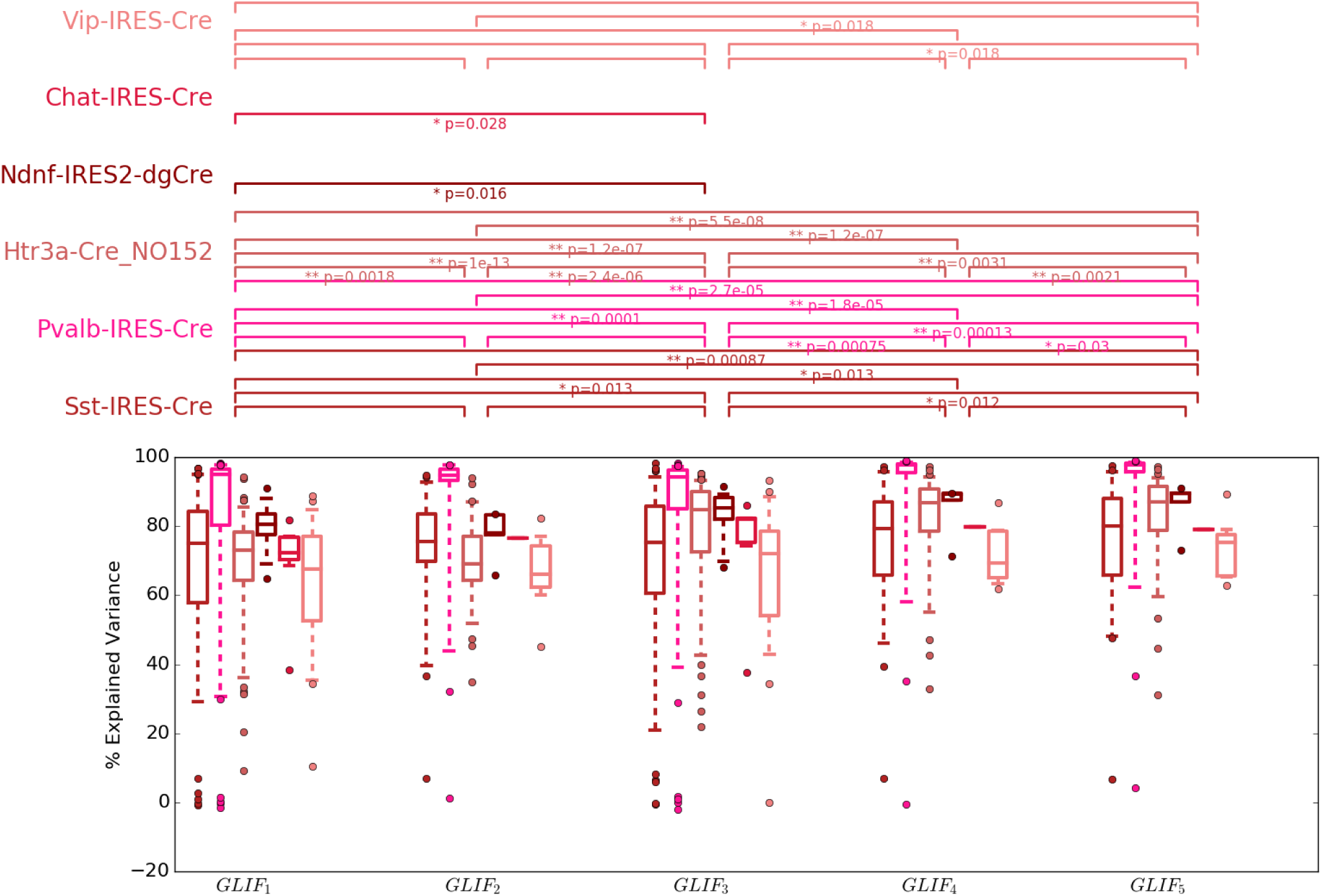
Explained variance summary for inhibitory GLIF models of different levels using threshold infinity obtained via MLIN optimization. Box plots display median, and quartiles. Whiskers reach to 5% and 95% quartiles. Individual data points lie outside whiskers. Brackets above plot denote signi cant difference between distributions assessed via a Wilcoxon sign ranked test: a single asterisk (*) represents a p-value < 0.05, a double asterisk (**) represents a p-value < 0.01. Distribution values are available in Table 4

**Figure 15:**
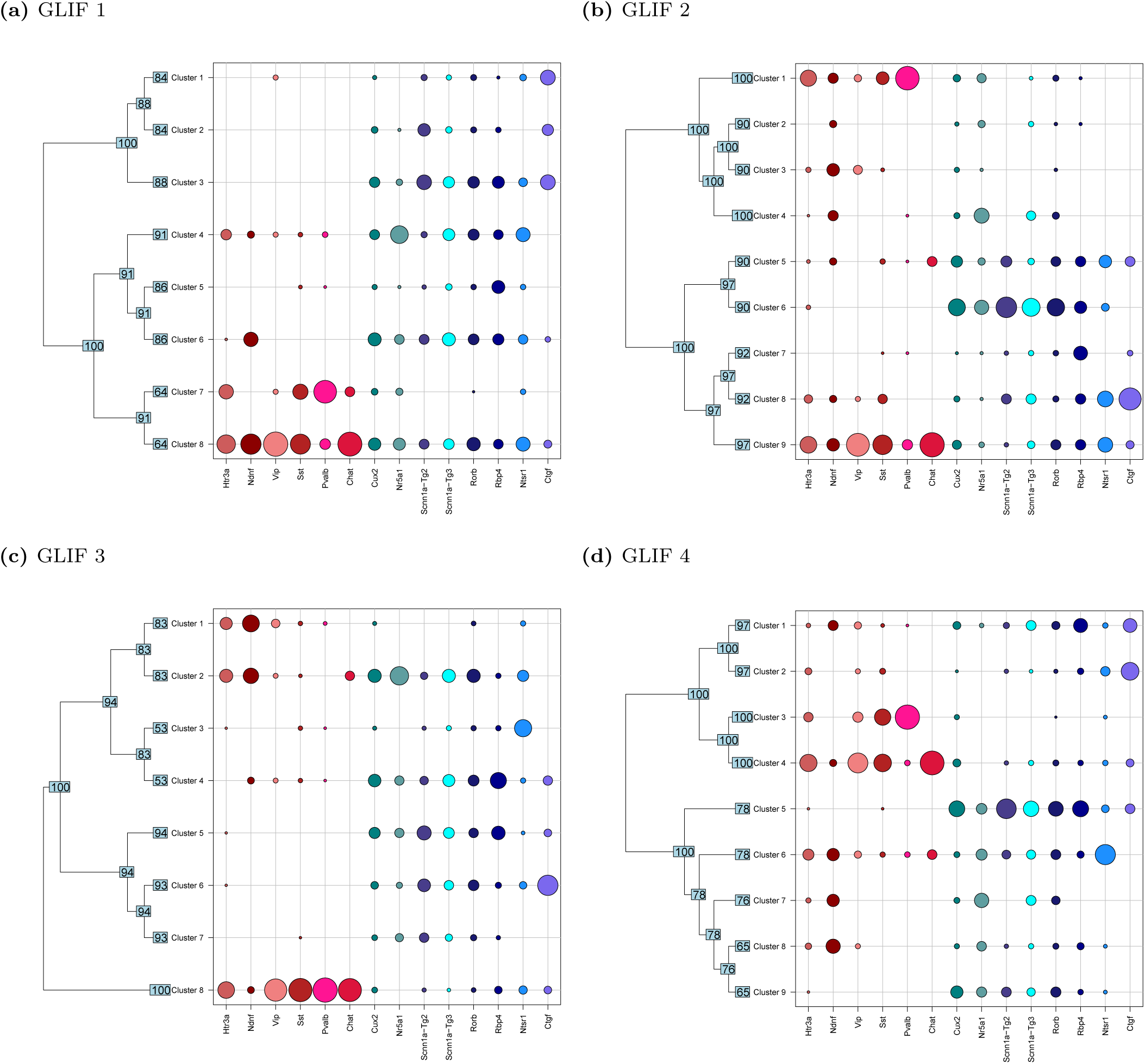
Clustering of cells using parameters obtained from four of the GLIF models described in the text (Table 1).

